# Tension-Tuned SynNotch Receptors for Synthetic Mechanotranduction and Intercellular Force Detection

**DOI:** 10.1101/2022.05.01.490205

**Authors:** D. Christopher Sloas, Jeremy C. Tran, Alexander M. Marzilli, John T. Ngo

## Abstract

Cells can sense and interpret mechanical stimuli from their environments and neighbors, but the ability to engineer customized mechanosensing capabilities has remained a synthetic and mechanobiology challenge. Here, we introduce tension-tuned synthetic Notch (SynNotch) receptors that can be used to convert extracellular and intercellular forces into specifiable gene expression changes. By elevating the tension requirements of SynNotch activation, in combination with structure-guided mutagenesis, we designed a set of receptors with mechanical sensitivities spanning the physiologically relevant picoNewton (pN) range. Cells expressing these receptors can distinguish between varying tensile forces and respond by enacting customizable transcriptional programs. The synthetic utility of these tools is demonstrated by designing a decision-making circuit, through which fibroblasts can be made to differentiate into myoblasts upon stimulation with distinct tension magnitudes. Mechanobiological utility is also demonstrated by characterizing cell-generated forces transmitted between cells during Notch signaling. Overall, this work provides insight regarding how mechanically induced changes in protein structure can be used to transduce physical forces into biochemical signals. The system should facilitate the further programming and dissection of force-related phenomena in biological systems.

## INTRODUCTION

Mechanical forces are fundamental regulators of biology, guiding vital processes and shaping disease progressions that span from the molecular to the tissue scale (*1–3*). Mechanical forces can do so by transmitting actionable information from biological environments, which can vary throughout developmental, tissue, and cell-cycle contexts (*4, 58*). Accordingly, cells have evolved complex mechanisms to actively utilize mechanical information, and to translate force-based stimuli into biochemical responses that can affect changes in metabolism, migration, differentiation, immunity, and gene transcription (*5–9*). Harnessing cells’ capability to interpret differential forces in their environment would enable advances in tissue engineering, where forces could be used to drive structural organization, as well as targeted therapeutics, where mechanical environments could be used as tractable disease biomarkers. However, it has been difficult to synthetically program mechanosensation in cells, given that many of its underlying mechanisms remain to be elucidated.

At a fundamental level, mechanotransduction can be generalized as an input-output relationship, in which mechanical energy is converted into a biochemical response via force-induced structural changes in cell signaling molecules. The sensitivity of proteins to mechanical unfolding has been tuned through several strategies, but in the context of biophysical spectroscopy, or FRET imaging sensors, rather than customized cellular signaling networks (*10–16*). Conversely, outputs from endogenous mechanotransduction pathways have been modulated by overexpressing protein components, engineering sensitivity to non-mechanical inputs, and, more synthetically, adding novel outputs to the force-sensitive Piezo1 and YAP/TAZ signaling cascades (*17–19*). However, these approaches have not yet tuned the mechanical force required for activation, and do not present an obvious protein engineering solution for doing so. These strategies furthermore rely upon endogenous pathways that are difficult to decouple from the user’s engineered pathways. Ideally, a fully customizable mechanosensitive pathway would synthesize three design criteria: 1) programmable sensitivity to input forces, 2) versatility of output cellular actions, and 3) orthogonality with respect to endogenous pathways.

Here, we use the Notch1 receptor to build a platform for controlling mechanosensation that meets the above design criteria. Notch1 is a classically studied protein, and our work capitalizes on a body of structural and biophysical research that guides our molecular design. Notch1 activation requires mechanical force, which is sensed by the receptor’s negative regulatory region (NRR) (**Fig. 1A**) (*20*). Upon application of tensile force via the ligand binding domain (LBD), the NRR undergoes a conformational change that enables proteolytic release of the intracellular domain (ICD), a transcriptional effector (*20–23*). Recent work has shown that the NRR can act as a modular scaffold to build synthetic Notch (SynNotch) receptors, which preserve the Notch activation mechanism while tolerating heterologous substitutions at the LBD and ICD (*24*). These receptors offer a starting point that satisfies two of our three design criteria; SynNotch receptors provide versatility of cellular outputs via an ICD of interest and minimal crosstalk with endogenous pathways via Notch’s mechanistically direct signaling pathway. We therefore asked if we could satisfy the remaining criterion—programmable sensitivity to input forces—by creating novel NRR domains with distinct mechanical properties, which could be used to specify new tension requirements for signaling.

**Fig. 1.**
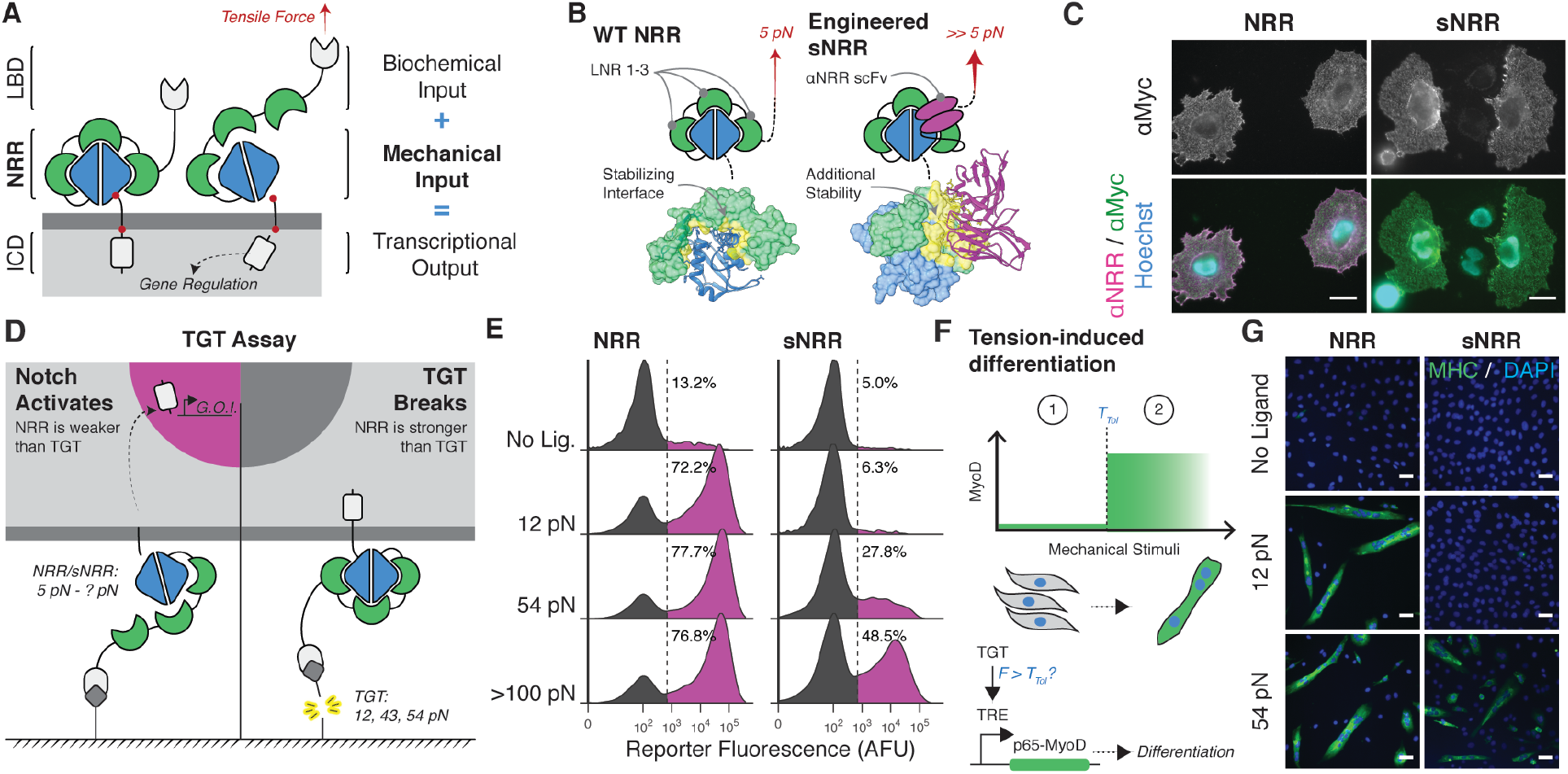
Design of customized mechanosensation. (**A**) Schematic of tension-mediated activation in Notch or SynNotch receptors. Application of sufficient tensile force via the LBD activates the receptor by displacing three LIN12-Notch repeat (LNR) modules and converting the NRR into a substrate for proteolysis at S2. Cleavage at S2, and concomitantly at S3, liberates the ICD. (LNR modules green; S2 and S3 red) (**B**) Activating the Notch1 NRR (left) requires tensile force to disrupt the intramolecular interactions that promote an autoinhibited conformation. Engineered sNRR domains (right) include an intramolecularly bound scFv for additional stability. The heterodimerization domain and LNR’s of the NRR are blue and green, respectively, and the scFv added in sNRR is magenta. Molecular interactions between the LNR’s and HD (left) or scFv and NRR (right) are yellow. (PDB 3ETO and 3L95, respectively) (**C**) Immunostaining of NRR- and sNRR-based receptors for surface-receptor with anti-myc (green) and available NRR with exogenous soluble anti-NRR scFv (magenta) in HeLa cells. Scale bar is 25 μm. (**D**) Schematic of TGT assay used to evaluate molecular tension needed to activate engineered receptors. Fluorescein (FITC) is used as a ligand for SynNotch receptors expressing an anti-FITC scFv LBD. (**E**) NRR- or sNRR-based receptors with a Gal4-VP64 ICD induce expression of a UAS-controlled H2B-mCherry reporter upon activation. HEK293FT cells transiently expressing these receptors were cultured on TGT-coated surfaces, and reporter activity was monitored by flow cytometry. (**F**) Tension-induced myogenic differentiation. C3H/10T1/2 fibroblasts express NRR- or sNRR-based SynNotch receptors with a tTA ICD. Upon activation, these receptors drive expression of p65-MyoD, which in turn leads to differentiation. (**G**) Immunostaining identifies differentiation into skeletal myocytes by multinucleation and positive myosin heavy chain (MHC) expression (green). Scale bar in C is 25 μm and 100 μm in G.

We began by designing a synthetic NRR domain that required increased tension levels to activate. Single molecule force spectroscopy has shown that proteins can be mechanically stabilized via the binding of other protein domains (*14, 25*), and studies of the NRR are consistent with these observations. Antibody binding can impart stability upon the NRR, elevating its resistance against chemical- and force-induced unfolding *in vitro* (*20, 26*) and limiting the spontaneous activation of mutant receptors *in vivo* (*27, 28*). Given these observations, we hypothesized that the NRR’s mechanical stability could be increased via direct fusion with an NRR-binding protein. Specifically, we fused the NRR with a single chain variable fragment (scFv) derived from an NRR-stabilizing antibody (*27*), such that the scFv could bind intramolecularly to the NRR to reinforce its autoinhibited conformation (**Fig. 1B**). In this way the integrated scFv can impart a mechanically stabilizing effect, and we anticipated that SynNotch receptors containing these “strengthened NRR” (sNRR) domains would require elevated tensions for activation. Here, we describe the development of tension-tuned sNRR-SynNotch receptors. We used these receptors to design force-sensitive genetic circuitries, including a mechanically activated cellular differentiation program. Further utility of the system is shown by their use in gauging the ligand pulling forces that are transmitted between cells during inter-cellular Notch signal transduction.

## RESULTS

### Synthetic mechanoreceptors are processed and activated similar to Notch

We first sought to verify that sNRR-based receptors preserved the fundamental trafficking and signaling functionalities of their NRR-based predecessors. To do so we compared receptors that differed only in the presence of a NRR or sNRR mechanosensing unit, having otherwise identical LBD and ICD components. As expected, both NRR- and sNRR-containing receptors were processed by the Golgi-localized furin convertase, and the resulting heterodimers were successfully trafficked to the cell surface (**Fig. 1C, S1A-B**). Labeling of live cells with soluble ligand indicated that the synthetic receptors were constitutively internalized from the plasma membrane (**Fig S1C**), similar to the known recycling of natural Notch receptors (*29*). Next, we demonstrated that integration of the anti-NRR scFv blocked sNRR reactivity against soluble anti-NRR, providing evidence that the fused antibody fragment interacts with the NRR epitope in the intended intramolecular manner (**Fig. 1C, S1B-C**). In addition to synthetic receptors, full-length Notch1 receptors maintained the desired expression properties upon substitution of the NRR for sNRR (**Fig. S1D-E**).

We used cell-based analyses with a luciferase reporter to confirm that sNRR-SynNotch could activate in response to ligand-mediated tension, as delivered through surface-immobilized fluorescein (FITC) dyes. FITC was chosen as a ligand due in part to its biocompatibility and versatility: it is a widely used research and clinical labeling reagent and thus diverse FITC-linked bioconjugates can be readily obtained. In addition, rupture forces between FITC and anti-FITC antibodies have previously been reported, with single molecule measurements indicating a ∼ 50 pN unbinding force between FITC and the E2 anti-FITC scFv (*57*). Given these factors, we paired the E2 anti-FITC scFv as a LBD for detecting FITC-based ligands.

Using cells expressing anti-FITC receptors, we observed that treatment with either batimastat (BB-94, a broad-spectrum metalloprotease inhibitor) or Compound E (a ɣ-secretase inhibitor) inhibited signaling responses against immobilized FITC ligands (**Fig. S2A**). This result indicates that sNRR activation involves proteolysis by a metalloproteinase and the ɣ-secretase complex, respectively, similar to natural Notch processing. The data also confirmed that sNRR-SynNotch could be activated upon exposure to an immobilized form of its ligand, suggesting a potential requirement for tensile resistance during activation, similar to natural Notch (*24, 30*). To verify this requirement, we treated cells with soluble fluorescein; such treatment failed to upregulate luciferase expression in both NRR- and sNRR-SynNotch cells (**Fig. S2B**). Thus, we conclude that sNRR-SynNotch processing is like that of natural Notch, and that sNRR-SynNotch also requires ligand-mediated tension for its signaling activity.

### Synthetic mechanoreceptors require elevated tension levels for signaling activity

Having confirmed its functional similarity to natural Notch, we next determined the force sensitivity of sNRR-SynNotch using the tension gauge tether (TGT) assay. In this approach, short double stranded DNA (dsDNA) sequences of known tension tolerance (T_tol_) are used to attach ligands (here, FITC) to a culture surface via biotin-streptavidin interaction prior to adding cells (**Fig. 1D**) (*21*). These dsDNA tethers (“TGTs”) impose a ceiling on the magnitude of tension that can be applied to individual ligand-bound receptors. If the tension needed to activate sNRR-SynNotch surpasses the T_tol_ of a given TGT, then the tether will dissociate prior to inducing receptor signaling. If instead the T_tol_ of a TGT surpasses what is needed to unravel the sNRR, then the tether is expected to endure and thus signaling should ensue. We used biotin connected to FITC via a non-rupturable linker, rather than dsDNA, as an upper limit for ligand T_tol_, as biotin-streptavidin will tolerate loads in excess of 100 pN prior to dissociation (*31*).

Previous studies have narrowed the activation force of the NRR to 4 – 12 pN (*20–23*), which is an order of magnitude below the critical force for rupturing antibody-antigen pairs (*32*). We therefore anticipated that sNRR would exhibit a significantly elevated force-activation threshold. By challenging receptor-expressing reporter cells against TGTs of distinct T_tol_ values, we defined an activation threshold of approximately > 54 pN for engineered sNRR-SynNotch receptors, well above the < 12 pN (∼ 5 pN) threshold previously defined for the Notch1 NRR (**Fig. 1E**). Other existing NRR-binding antibodies were fused as scFv’s to the NRR to evaluate the generalizability of our design, and multiple increased receptor tension tolerance (*28, 33*) (**Fig. S3**). These data demonstrate that sNRR fusions have increased force requirements for activation since they respond differentially to molecular tensions even while binding the same ligand.

### Synthetic mechanotransduction can induce muscle-cell like phenotypes

Equipped with two mechanically distinct receptors, we next aimed to specify how a cell responds to a specific force value. In natural systems, cells enact intricate biological outputs based upon mechanical information in their microenvironment. For example, mechanical forces can guide formation of sarcomeres (*34*), coordinate wound healing (*35*), or induce differentiation (*7*). Using a synthetic biology approach, we generated mouse embryonic fibroblasts that can differentiate in response to specific molecular tensions. To do so, we rendered MyoD, a master regulator of myogenic differentiation, SynNotch-dependent. A p65-MyoD fusion under control of the *tetO*-containing TRE promoter (TRE: p65-MyoD) was virally integrated into C3H/10T1/2 fibroblasts (*36*). Receptors containing *tetO*-binding ICDs (TetR-VP64) were then expressed in these cells, such that signaling would lead to p65-MyoD expression and subsequently myogenic conversion (**Fig. 1F**). Using these engineered fibroblasts, we then compared the ability of NRR-SynNotch and sNRR-SynNotch to guide cell fate decisions in response to mechanically distinct TGTs. We observed that cells expressing the receptors differentiated and fused into multinucleated skeletal myocytes only in response to tensions greater than the threshold defined by their respective mechanosensing domains—at least 12 pN for the NRR, and at least 54 pN for sNRR (**Fig. 1G, S4**).

### Receptor tensile strengths can be systematically tuned by mutation

Further intricacy in cellular mechanosensation can arise from protein isoforms that have distinct mechanical properties despite overlapping biochemical functions, as studied with integrins or talins (*37, 38*). We therefore anticipated that an expanded family of sNRR domains with diversified force activation thresholds would similarly enable more sophisticated signaling networks. Mutating residues involved in non-covalent interactions at a protein’s “mechano-active” site can lead to distinct mechanical phenotypes (*39*). We hypothesized that point mutations within the scFv:NRR interface would alter the mechanical stability of sNRR, providing a basis to systematically tune force activation thresholds (**Fig. 2A**). Using a structure-guided strategy, we generated single- and double-mutants by introducing conservative and non-conservative amino acid substitutions within the scFv:NRR interface (**Fig. 2B-C**).

**Fig. 2.**
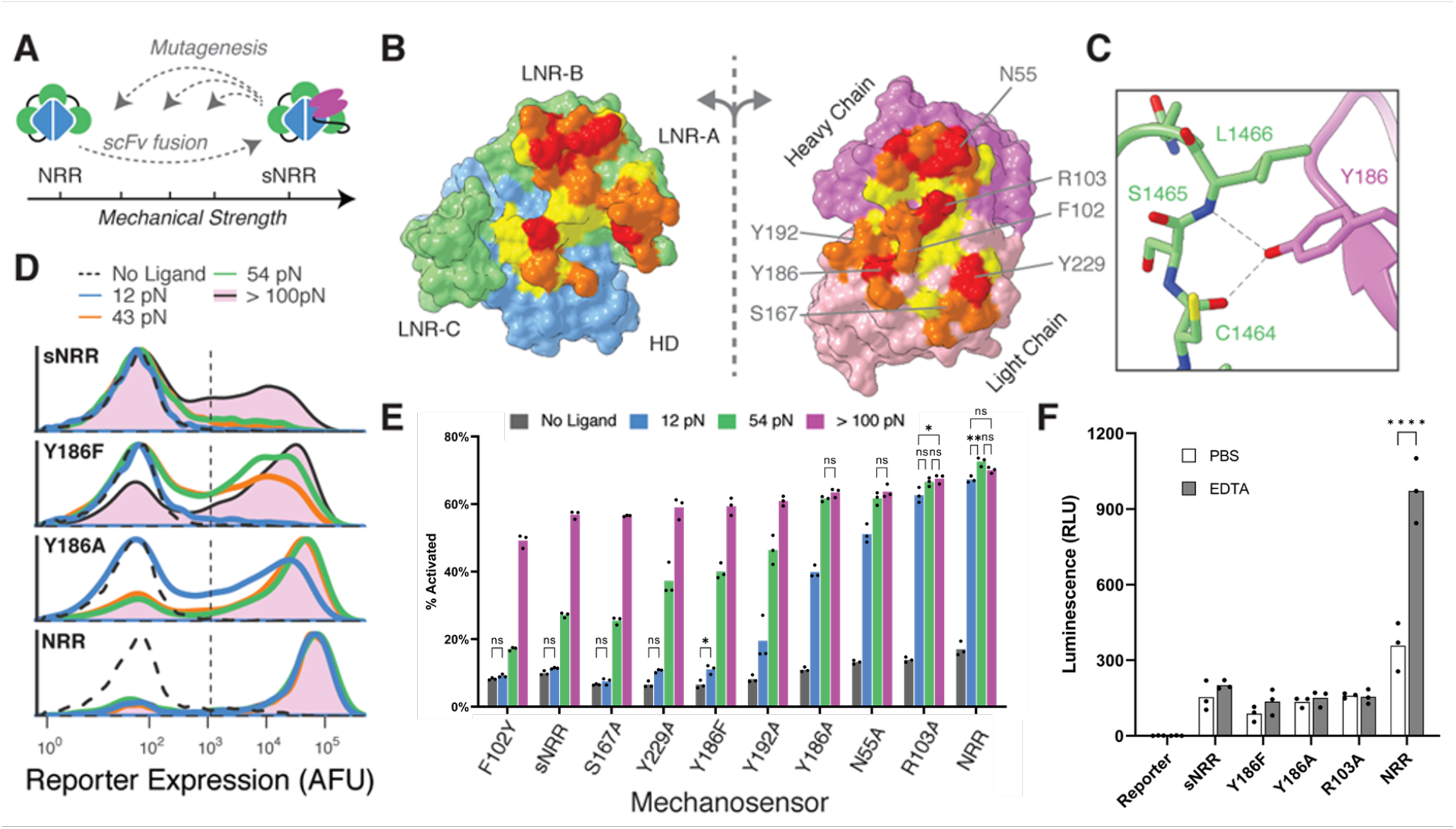
Tuning the tensile strength of sNRR domains. (**A**) Schematic of strategy to tune mechanical strength. (**B**) Open-book view of the interface between the Notch1 NRR and the NRR-binding Fab. The HD and LNR’s of the NRR are blue and green, respectively, and the heavy and light chains of the scFv are magenta and pink. Additional residue coloring corresponds to the distance between the two surfaces (< 5.5 A, yellow; 5.5 – 4 A, orange; 4 – 3 A, red) (PDB 3L95). (**C**) Y186 on the Fab light chain is highlighted as an example of a mutated residue. (**D**) Mutating Y186 decreases sNRR mechanical strength. Receptors stably integrated in HEK293FT cells upregulate UAS-driven H2B-mCherry reporter upon activation with TGTs. (**E**) Mutating sNRR residues creates receptors with a range of force activation thresholds. Receptors are transiently expressed in HEK293FT cells with stably integrated UAS-H2B-mCherry reporter. (**F**) Cells expressing SynNotch receptors with an NRR domain or sNRR domain of various strengths are treated with either 0.5 mM EDTA or PBS for 15 minutes at 37 °C, then treatment was quenched with culture media. Reporter expression was evaluated 6 hours later. Activated receptors upregulate expression of a UAS-regulated luciferase. Biological triplicate, 2-way ANOVA where null hypotheses cannot be rejected, labeled n.s. (f > 0.05), * (f < 0.05), or unlabeled and (f < 0.001) for each receptor unlabeled comparisons.

Consistent with our expectation, introducing mutations resulted in sNRR units of distinct mechanical strengths, as quantified using a fluorescent reporter gene and the TGT assay (**Fig. 2D-E**). Mutation generally led to sNRR isoforms with lower activation thresholds, resulting in domains with varying tension tolerances against biologically relevant pN levels (*40*) and allowing us to populate the sensitivity gap separating NRR and sNRR domains. For example, conservatively mutating Tyr186 to Phe decreased the activation threshold below 54 pN, and more severely mutating to Ala furthered this effect. Weakening single mutations were generally additive when combined into doubly mutated domains (**Fig. S5A**). One mutation (F102Y) may have led to an increase in mechanical stability, though marginal and difficult to resolve using TGTs (**Fig. S5B-C**).

Of particular use was the Y186F mutation (sNRR_Y186F_), which is weakened such that it activates to a greater extent than the original sNRR on strong TGTs while still resisting activation by 12 pN TGTs. Immunostaining confirmed that mutated receptors were trafficked to the cell surface with comparable efficiencies (**Fig. S6**). We also note that sNRR-containing receptors exhibit reduced ligand-independent activation (LIA) in the absence of stimulus, as compared to their NRR-based counterpart (**Fig. 2E**). Reduced LIA has been targeted as a means to improve SynNotch functionality (*41*). The difference in LIA for sNRR is increasingly apparent when the ICD is replaced with more potent transcriptional activators VP64-p65 or VP64-p65-RTA (VPR) (**Fig. S7**).

Further analysis of sNRR mechanical strength showed that scFv insertion provides the primary mechanism of stabilizing these engineered domains. In T-cell acute lymphoblastic leukemia (T-ALL), mutations within the NRR can result in destabilized domains that readily undergo LIA (*42*). However, these same T-ALL mutations, when distal to the scFv binding site, did not affect the mechanical strength of sNRR. Instead, presence of the scFv rescues the hyperactive phenotype, providing evidence that scFv insertion provides stabilization that is a new rate-limiting step in receptor activation (**Fig. S8**). Only NRR mutations at the scFv binding interface destabilized sNRR-SynNotch receptors, consistent with results from our collection of scFv mutations. NRR mechanical stability depends on the presence of calcium ions, and LIA can be ectopically induced through calcium chelation using EDTA (*43*). Interestingly, sNRR domains resist activation by EDTA treatment (**Fig. 2F**). This result holds true for mechanically weakened sNRR variants, including the highly destabilized sNRR_R103A_ mutant, which was similar to NRR-SynNotch in TGT analyses. Together, these results indicate that scFv fusion provides a dominant autoinhibitory effect. As EDTA is a routine reagent used in cell culture, sNRR domains serendipitously provide a SynNotch variant that can be stably integrated and expanded in cells while resisting inadvertent LIA.

### Genetic circuits can specify distinct gene expression responses in cells experiencing differing tension levels

Using our set of sNRR domains, we demonstrated further versatility in cellular outputs by designing gene circuits that selectively filter various magnitudes of tension. Synthetic and natural signaling networks can use filtering logic to detect an input, such as small molecule concentration, only if it falls within a desired range. Described in this framework, the mechanogenetic circuits shown thus far generate “high-pass filters,” in which a target gene is expressed only in response to tension beyond a threshold T_tol_. Recognizing the engineering utility of “low-pass” and “band-pass” filters that detect low and intermediate forces, respectively, we sought to extend our sNRR-based genetic circuits to achieve these complex outputs.

To build a low-pass filter, we devised a circuit in which a target gene is expressed only if tensions fall below an activation threshold (**Fig. S9A**). In this approach, activation of a sNRR_Y186F_-SynNotch receptor directs the transcription of a microRNA (miRNA), which in turn inhibits the translation of a constitutively transcribed fluorescent protein gene based on mCerulean (CMV-mCer-target-miR-FF4, see Methods) (**Fig. S9B**). We observe that this circuit downregulates mCerulean fluorescence in response to increasing molecular tension, as designed. This behavior is reminiscent of natural mechanisms in which mesenchymal stem cells use miRNA to downregulate gene expression in response to stiff ECM (*44*). In addition to the transcriptional systems demonstrated thus far, the use of miRNA also highlights the ability to install force-activated post-transcriptional control.

In addition to a low-pass filter, we also engineered a band-pass capable of detecting intermediate force magnitudes. Here, we reasoned that band-pass behavior could more closely mimic that of induced pluripotent stem cells (iPSCs), which pursue distinct cell fates in response to soft, medium, or rigid ECM stiffnesses (*7*). To implement such logic, we combined multiple receptors in a single cell to implement an incoherent feedforward loop (IFFL) (**Fig. 3A**). The IFFL is a signaling motif commonly employed to create a band-pass response with respect to time or concentration (*45, 46*), and we hypothesized that we could extend the IFFL to similarly create a band-pass behavior with respect to molecular tension. In the activating arm of the loop, an NRR-SynNotch possessing a Gal4-VP64 ICD is used to drive expression of a cyan fluorescent mCerulean reporter (UAS-mCer). In the repressing arm of the loop, a sNRR_Y186F_-SynNotch with a TetR-based ICD is used to drive a red fluorescent Gal4-KRAB fusion, which is able to repress UAS-regulated mCerulean expression. Co-transfected cells encoding the IFFL circuit exhibited the expected response profiles, with mCerulean expression occurring only in response to intermediate tension values (**Fig. 3B-D**).

**Fig. 3.**
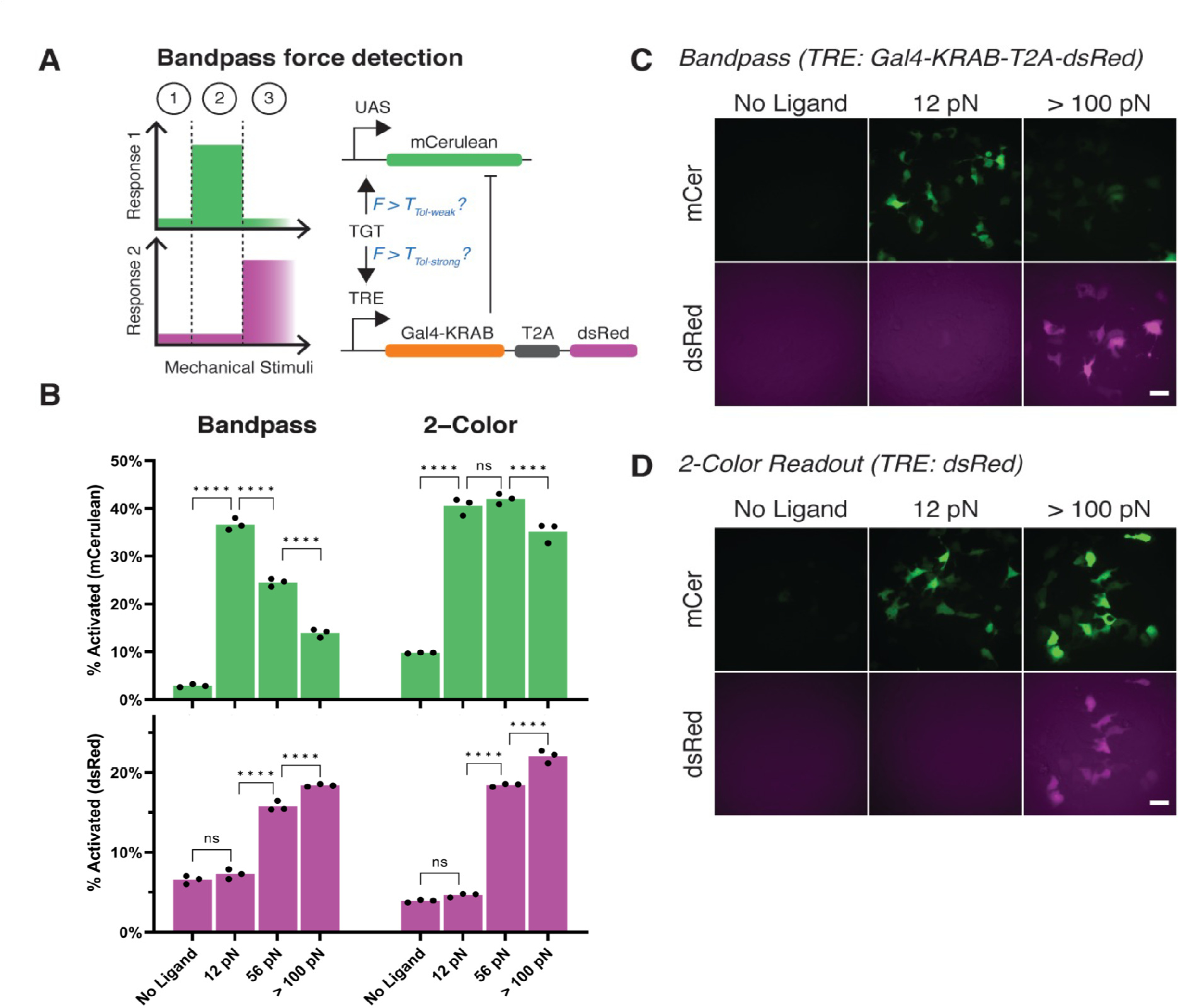
Bandpass filtering in mechanogenetic circuits. (**A**) An IFFL is designed to respond to intermediate forces. A NRR-based SynNotch with Gal4-VP64 ICD drives mCerulean (mCer) reporter activity upon activation. A sNRR_Y186F_-based SynNotch with tTA ICD drives DsRed-Express2 (dsRed) expression in combination with Gal4-KRAB to inhibit mCer production. (**B**) mCer (top) and dsRed (bottom) activation of mechanogenetic circuits. “Bandpass” (left) denotes cells expressing the full bandpass circuit in (A). The “2-Color” designation (right) denotes cells expressing this circuit without the Gal4-KRAB inhibitory component. (**C-D**) Fluorescence images of reporter activity in HEK293-FT cells expressing Bandpass or 2-Color readout circuits. Scale bars are 25 μm.

We also tested a circuit in which sNRR_Y186F_-SynNotch was made to activate a TRE-DsRed-Express2 (TRE-DsRed-E2) reporter, without repressing UAS-mCerulean. As expected, cells grown on high tension ligands expressed both mCerulean and DsRed-E2 reporters, whereas intermediate tension ligands activated only mCerulean (**Fig. 3D**). This result confirms the IFFL circuit function and shows that cells can be made to express distinct gene combinations in response to different ligand tension levels. In this case, dual expression of cyan and red fluorescent proteins provides a “2-color” readout that is indicative of cell engagement with high tension ligands.

### Force transmission depends on ligand-receptor bond properties

Having demonstrated their synthetic utility, we next aimed to apply our receptors to measure inter-cellular force transmissions. To do so, we replaced TGTs with encodable ligands based on GFP, which was selected given its ability to interact with the LaG17 nanobody present within our receptor ECDs. Prior to testing cell-based ligands, we determined whether LaG17-GFP complexes could endure the tension levels required for sNRR activation using streptavidin-immobilized GFP. sNRR-SynNotch cells were refractory under these conditions, suggesting a restriction in the tension levels that can be propagated via LaG17-GFP bonds. To overcome this limitation, LaG17 was replaced with the LaG16 nanobody, which we expected to bear increased tension loads against GFP, given its reduced off-rate constant (k_off_) compared to that of LaG17 (k_off_ = 1.1 × 10^−3^ s^-1^ for LaG16-GFP; k_off_ = 1.4 × 10^−1^ s^-1^ for LaG17-GFP) (*47*). Consistent with this prediction, LaG16-mediated binding resulted in strong sNRR-SynNotch signaling against immobilized GFP (**Fig. S10**). We thus proceeded with LaG16-containing receptors in *trans*-activation analyses.

### Intercellular pulling forces can be gauged via cell-based trans-activation

In natural systems, the mechanical pulling activity of Notch ligands requires their intracellular ubiquitination and entry into the epsin-dependent clathrin mediated endocytosis (CME) pathway (*48*). Mammalian Notch ligands are ubiquitinated by the E3 ligase Mindbomb-1 (MIB-1) then bound by epsins, which facilitate clathrin-mediated endocytosis (CME) (*49*). During CME, epsins couple with polymerizing actin filaments (*50*) and physically link invaginating pits to cellular force-generating machinery (*51-52*). Although many of the factors required for epsin-dependent uptake are known (*53*), the pulling capacity that they confer to ligands during CME is unknown. To probe this capacity, we next generated a synthetic GFP-based ligand by fusing GFP to the transmembrane domain (TMD) and intracellular tail of the native Notch ligand DLL1 (**Fig. 4A**, GFP-TMD-DLL1).

**Fig. 4.**
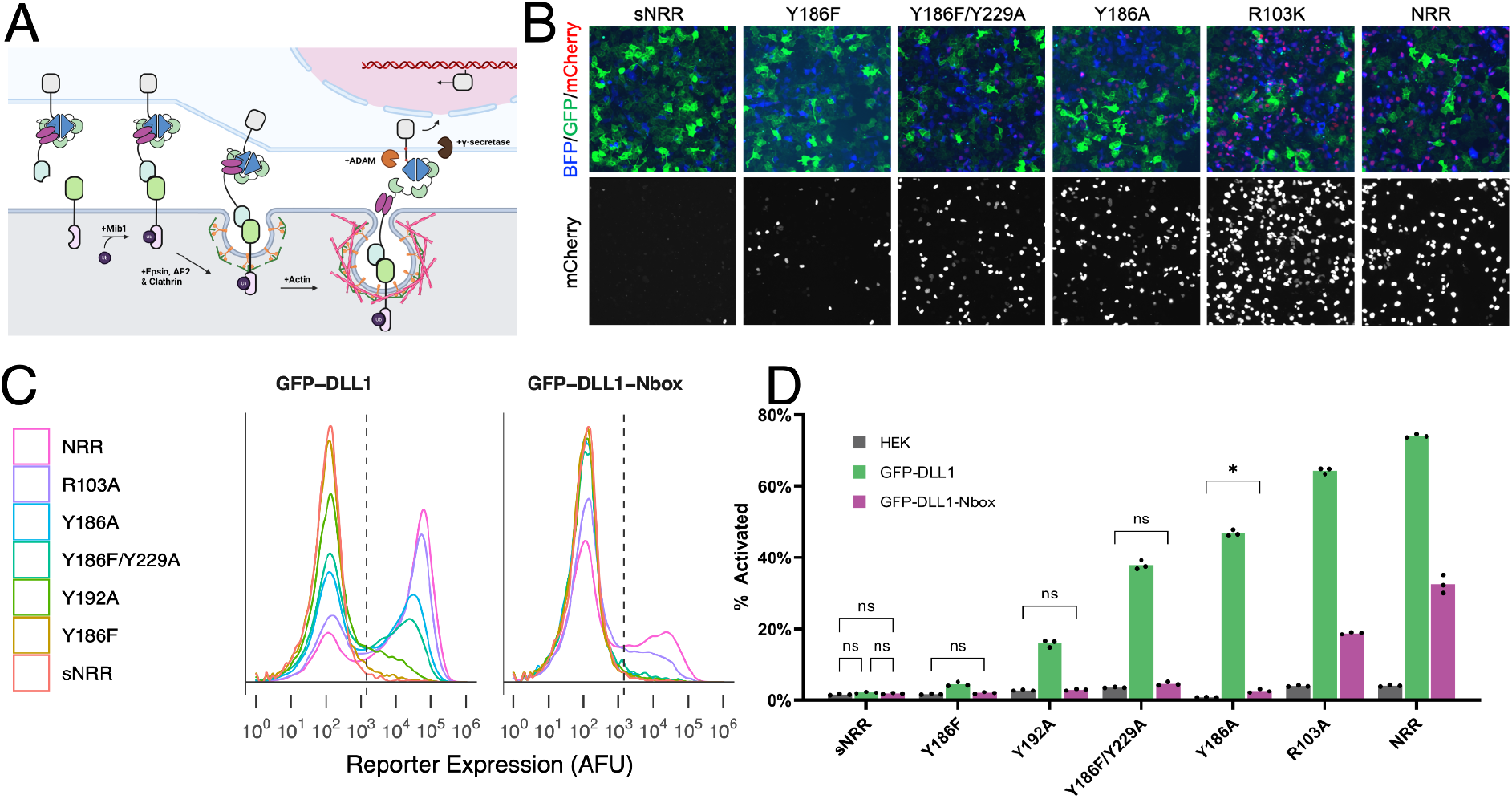
Intercellular mechanotransduction via cell-generated, ubiquitination-dependent endocytic forces. (**A**) Schematic of ligand-CME transactivating Notch receptors. DLL1 ligand uptake is regulated by Mib-1 ubiquitination, which is recognized by cargo-specific adaptor proteins (epsins) that link ligands to CME machinery. Endocytosis generates a pulling force which exposes the NRR’s cryptic S2 proteolytic site, activating the receptor. (**B**) LaG16-sNRR variants are *trans*-activated by GFP-TMD-DLL1 senders to varying degrees. Receptors stably integrated in HEK293-FT cells upregulate UAS-driven H2B-mCherry reporter upon activation by HEK293-FT GFP-TMD-DLL1 in 24-hour coculture. Inset is H2B-mCherry channel under increased contrast. (**C**) Flow cytometry traces of receiver cell reporter expression levels following coculture with either GFP-TMD-DLL1 or GFP-TMD-DLL1-Nbox sender cells. Traces represent normalized densities of receiver (BFP+) cells within the analyzed coculture populations. Dashed vertical lines indicate H2B-mCherry+ threshold fluorescence values. (**D**) Percent of H2B-mCherry+ reporter cells in receiver cell population (BFP+) from (B), along with signals from control cocultures using HEK293-FT mock ligand cells. Biological triplicate, 2-way ANOVA where null hypotheses cannot be rejected, labeled n.s. (f > 0.05), * (f < 0.05), or unlabeled (f < 0.001) for each receptor; > 50,000 cells analyzed per data replicate.

GFP-TMD-DLL1 was stably expressed in HEK293-FT cells to generate ligand-expressing “sender” cells. Senders expressing a MIB1-resistant (and thus epsin-binding deficient) ligand were also generated, by expressing a mutant ligand containing an ablated “N-box” motif (GFP-TMD-DLL1-Nbox), required for MIB1 binding (*49, 54*). Live cell immunostaining confirmed the presentation of these ligands on cell surfaces at comparable levels (**Fig. S11**). To evaluate their activities, we co-cultured sender cells with receptor-expressing “receiver” cells, allowing *trans*-cellular stimulation to proceed for 24 hours prior to analyzing receiver cell reporter levels. These analyses confirmed the mechanical activity of GFP-TMD-DLL1 senders, which *trans*-activated receivers expressing multiple distinct mechanoreceptor domains (**Fig. 4B-D**). Quantification of signaling intensities showed agreement with TGT-based measurements: domains that resisted 12 pN tethers (including sNRR and sNRR_Y186F_) were refractory against GFP-TMD-DLL1, whereas those that with sensitivity to 12 pN tethers were similarly susceptible against GFP-TMD-DLL1 in *trans*.

Comparison of signaling levels induced by GFP-TMD-DLL1 versus GFP-TMD-DLL1-Nbox senders revealed CME-dependent and -independent activities. Receptors containing mechanically weak domains (NRR and sNRR_103A_) were activated at partial levels due to cell-cell contact with GFP-TMD-DLL1-Nbox sender cells (**Fig. 4 C-D**). This result suggests that lateral forces due to trans-cellular LaG16-GFP binding, may supply sufficient forces to trigger signaling independent of CME. In contrast, receptors with intermediate strength domains (including sNRR_Y186A_ and sNRR_Y192A_) showed explicit requirements for CME, whereas high stability receptors (including sNRR and sNRR_Y186F_) largely resisted CME-generated tensions. Of note, the CME-resistant sNRR_Y186F_ domain could be rendered CME-susceptible following addition of a second destabilizing mutation (sNRR_Y186F/Y229A_). These data indicate that ligand uptake, regulated through MIB1-mediated ubiquitination, can confer tension at levels beyond the NRR’s activation threshold (∼ 5 pN), yet below what is needed for sNRR_Y186F_ (> 12 pN). Overall, our results confirm that sNRR-based receptors can detect and distinguish between pN levels of intercellular forces.

### Bifunctional bridges can induce mechanical coupling between cells

Recognizing that mechanical signaling events are highly coordinated *in vivo*, we next sought to gain spatiotemporal control over intercellular force transmissions. To do so, we implemented an inducible mechanical coupling strategy using a bispecific small molecule bridge. In this approach, we used a ligand that contains a biotinamide-binding antibody fragment (*55-56*) followed by a transmembrane domain and the DLL1 intracellular tail (anti-biotin-TMD-DLL1). Tests using immobilized biotin ligands showed that biotin:anti-biotin bonds could withstand the tension levels needed for sNRR activation (**Fig. S12**). Furthermore, *trans*-activation between anti-biotin-TMD-DLL1 senders and anti-FITC receivers could be induced via treatment with soluble biotin-FITC at sub-nanomolar concentrations (**Fig. S13, Fig. S14D, Supplementary Movie 1**). To quantify drug-induced signaling levels we used U2OS receiver cells containing a UAS-DsRed-Express2 reporter gene. Receivers were mixed with anti-biotin-TMD-DLL1 senders and reporter levels were quantified after overnight treatment with 2 nM biotin-FITC. Under these conditions, receiver cells activity levels were consistent with our TGT- and GFP-TMD-DLL1-based measurements, and the endocytic requirement of these signals was confirmed via co-treatment with the dynamin inhibitor dynasore (**Fig. S14B**). Together, these results further confirm the distinct mechano-sensitivities of our tension-tuned receptors.

Finally, to achieve spatiotemporal control, we applied removable microwell inserts to generate defined sender-receiver boundaries (**Fig. S14C**). Following microwell removal and cell growth, biotin-FITC was added to cultures to activate coupling between sender and receiver cells. After overnight incubation with the drug, fluorescence imaging revealed spatially confined signaling activities that were limited to receiver cells residing at sender-receiver boundaries (**Fig. S14D**). These results confirmed the requirement of sender-receiver contact in generating biotin-FITC induced signaling responses.

## DISCUSSION

In summary, we developed tension-tuned SynNotch receptors that can serve as genetically encoded tensiometers. These receptors can be used not only to engineer mechanical signaling responses, but also to decipher and record tensile forces as quantifiable readouts of a cell’s current or past mechanical interactions. The versatility of the system enabled force detection via diverse ligand forms, including FITC-labeled TGTs, GFP-tagged proteins, and a bifunctional small molecule based on biotin-FITC.

To permit the tuning of receptor sensitivities, we first stabilized the NRR by fusing it with an inhibitory anti-NRR antibody fragment, producing a strengthened NRR (sNRR) that required ten-times more tension for signaling activity. This requirement was then lowered by altering binding interactions within the domain’s “mechanoactive” site using a structure-guided mutagenesis approach. This strategy allowed us to populate the sensitivity gap between NRR and sNRR domains, resulting in 14 mechanoreceptor sequences with tensile sensitivities that span the biologically relevant pN range. When expressed by cells, these proteins could be used to activate gene expression responses following stimulation with extracellular and intercellular tensional cues.

We demonstrated the utility of our system in two important ways: first, in a synthetic biology approach, we applied our receptors to design cells that could distinguish between mechanical signals and enact customized transcriptional responses in turn. This strategy was used to implement mechanical “filtering logic,” which enabled cells to express a reporter gene only in response to intermediate tension values. A tension-dependent differentiation circuit was also developed, and we used this program to define the force levels needed to induce muscle cell-like phenotypes in fibroblast-based progenitor cells.

In a biological application, we used our receptors to measure cell-generated, biomechanical forces exchanged between cells during Notch signaling. We showed that chimeric ligands containing DLL1 intracellular tails could deliver tensions at levels beyond the ∼5 pN rupture requirement of the NRR, yet at levels below that which is supplied by 12 pN TGTs. These values agree with previous laser-trapping measurements, where DLL1 cells were found to pull Notch-coated beads with ∼ 10 pN energies (*60*). Like native DLL1, the pulling capability of chimeric ligands also required MIB1-binding and dynamin activity.

There should be several advantageous applications of our tools moving forward. For example, intermediate-strength receptors (which explicitly required ligand CME for signaling) could be used in mapping synaptic connections between neurons by exploiting pre-synaptic forces due to recycling neurotransmitter vesicles. Such an approach may provide a specific and encodable anterograde tracing method in which signaling due to non-synaptic cell-cell proximity could be overcome. Furthermore, combined use with drug-based bridging agents could be used to achieve temporal resolution in connectivity mapping, which would allow researchers to monitor changes in the formation and elimination of synapses over time.

During our experimentation, we identified a design feature that should be considered in future adaptations of the method—that ligand-receptor bonds must be of sufficient tensile strength in order to propagate forces to NRR/sNRR domains. Thus, new ligand-receptor pairings should be tested using immobilized ligands prior to *trans*-cellular analyses. Antibody-antigen interactions are recommended as a starting point for new designs, given that these complexes generally form strong bonds (*32*). Antibody-antigen bonds could be directly encoded as ligand-receptor pairings or exploited by combining anti-FITC receptors with FITC-conjugated antibodies or polypeptides as soluble bridging agents. Such an approach could be used to investigate mechanosensitive pathways beyond Notch signaling, including those involving integrins, cadherins, and immune cell antigen receptors.

Finally, our work demonstrates how biological insights regarding natural mechanisms can guide the design of sophisticated and useful cell signaling components. The data also reinforce a long-established paradigm in protein engineering: that by adding extra stability, one can render proteins more tolerant to mutation, and thus more susceptible to engineering (*59*). Although our work focused on transcription-based outputs, future adaptations of the method could enable direct mechanical control over processes such as genome editing, metabolism, or cell-killing activity. Use of iterative or continuous library screening methods may accelerate the development of such systems. As mechanosensitive proteins continue to be structurally and mechanistically elucidated, we anticipate that the mechanogenetic “toolkit” will continue to grow; such expansion should in turn facilitate development of novel targeted therapies and tissue-engineered systems.

## Supporting information

Supplementary Information for v2

## ACKNOWLEDGMENTS

D.C.S. and A.M.M. were supported through Graduate Research Fellowship awards from the National Science Foundation. J.C.T was supported through a Cross-Disciplinary Fellowship awarded through BUnano (Boston University Nanotechnology Innovation Center). Funding for this work was provided through NIH research grants from the NIGMS (R35 GM128859) and the NHLBI (R01 HL147585).

## REFERENCES

1. I. Schoen, B. L. Pruitt, V. Vogel, The Yin-Yang of Rigidity Sensing: How Forces and Mechanical Properties Regulate the Cellular Response to Materials. Annu. Rev. Mater. Res. 43, 589–618 (2013).

2. R. G. Wells, Tissue mechanics and fibrosis. Biochim. Biophys. Acta - Mol. Basis Dis. 1832 (2013), pp. 884–890.

3. C. Hahn, M. A. Schwartz, Mechanotransduction in vascular physiology and atherogenesis. Nat. Rev. Mol. Cell Biol. 10, 53–62 (2009).

4. T. Mammoto, A. Mammoto, D. E. Ingber, Mechanobiology and Developmental Control. Annu. Rev. Cell Dev. Biol. 29, 27–61 (2013).

5. R. Bertolio, F. Napoletano, M. Mano, S. Maurer-Stroh, M. Fantuz, A. Zannini, S. Bicciato, G. Sorrentino, G. Del Sal, Sterol regulatory element binding protein 1 couples mechanical cues and lipid metabolism. Nat. Commun. 10, 1326 (2019).

6. S. Van Helvert, C. Storm, P. Friedl, Mechanoreciprocity in cell migration. Nat. Cell Biol. 20, 8–20 (2018).

7. A. J. Engler, S. Sen, H. L. Sweeney, D. E. Discher, Matrix Elasticity Directs Stem Cell Lineage Specification. Cell. 126, 677–689 (2006).

8. D. K. Das, Y. Feng, R. J. Mallis, X. Li, D. B. Keskin, R. E. Hussey, S. K. Brady, J.-H. Wang, G. Wagner, E. L. Reinherz, M. J. Lang, Force-dependent transition in the T-cell receptor β-subunit allosterically regulates peptide discrimination and pMHC bond lifetime. Proc. Natl. Acad. Sci. U. S. A. 112, 1517–1522 (2015).

9. A. Elosegui-Artola, I. Andreu, A. E. M. Beedle, A. Lezamiz, M. Uroz, A. J. Kosmalska, R. Oria, J. Z. Kechagia, P. Rico-Lastres, A. L. Le Roux, C. M. Shanahan, X. Trepat, D. Navajas, S. Garcia-Manyes, P. Roca-Cusachs, Force Triggers YAP Nuclear Entry by Regulating Transport across Nuclear Pores. Cell. 171, 1397-1410.e14 (2017).

10. D. Sharma, O. Perisic, Q. Peng, Y. Cao, C. Lam, H. Lu, H. Li, Single-molecule force spectroscopy reveals a mechanically stable protein fold and the rational tuning of its mechanical stability. Proc. Natl. Acad. Sci. U. S. A. 104, 9278–9283 (2007).

11. T. Bornschlögl, J. Christof M Gebhardt, M. Rief, Designing the folding mechanics of coiled coils. ChemPhysChem. 10, 2800–2804 (2009).

12. S. P. Ng, K. S. Billings, T. Ohashi, M. D. Allen, R. B. Best, L. G. Randles, H. P. Erickson, J. Clarke, Designing an extracellular matrix protein with enhanced mechanical stability. Proc. Natl. Acad. Sci. U. S. A. 104, 9633–7 (2007).

13. D. P. Sadler, E. Petrik, Y. Taniguchi, J. R. Pullen, M. Kawakami, S. E. Radford, D. J. Brockwell, Identification of a Mechanical Rheostat in the Hydrophobic Core of Protein L. J. Mol. Biol. 393, 237–248 (2009).

14. Y. Cao, T. Yoo, S. Zhuang, H. Li, Protein-Protein Interaction Regulates Proteins’ Mechanical Stability. J. Mol. Biol. 378, 1132–1141 (2008).

15. Y. Cao, T. Yoo, H. Li, Single molecule force spectroscopy reveals engineered metal chelation is a general approach to enhance mechanical stability of proteins. Proc. Natl. Acad. Sci. 105, 11152–11157 (2008).

16. P. Ringer, A. Weißl, A.-L. Cost, A. Freikamp, B. Sabass, A. Mehlich, M. Tramier, M. Rief, C. Grashoff, Multiplexing molecular tension sensors reveals piconewton force gradient across talin-1. Nat. Methods. 14, 1090–1096 (2017).

17. J. H. Hughes, S. Kumar, Synthetic mechanobiology: Engineering cellular force generation and signaling. Curr. Opin. Biotechnol. 40 (2016), pp. 82–89.

18. Y. Pan, S. Yoon, J. Sun, Z. Huang, C. Lee, M. Allen, Y. Wu, Y.-J. Chang, M. Sadelain, K. K. Shung, S. Chien, Y. Wang, Mechanogenetics for the remote and noninvasive control of cancer immunotherapy. Proc. Natl. Acad. Sci., 201714900 (2018).

19. L. N. Liu, S. X. Zhang, W. B. Liao, H. P. Farhoodi, C. W. Wong, C. C. Chen, A. I. Ségaliny, J. V. Chacko, L. P. Nguyen, M. R. Lu, G. Polovin, E. J. Pone, T. L. Downing, D. A. Lawson, M. A. Digman, W. A. Zhao, A. I. Segaliny, J. V. Chacko, L. P. Nguyen, M. R. Lu, G. Polovin, E. J. Pone, T. L. Downing, D. A. Lawson, M. A. Digman, W. A. Zhao, Mechanoresponsive stem cells to target cancer metastases through biophysical cues. Sci. Transl. Med. 9 (2017)

20. W. R. Gordon, B. Zimmerman, L. He, L. J. Miles, J. Huang, K. Tiyanont, D. G. McArthur, J. C. Aster, N. Perrimon, J. J. Loparo, S. C. Blacklow, Mechanical Allostery: Evidence for a Force Requirement in the Proteolytic Activation of Notch. Dev. Cell. 33, 729–736 (2015).

21. X. Wang, T. Ha, Defining Single Molecular Forces Required to Activate Integrin and Notch Signaling. Science, 340, 991–994 (2013).

22. F. Chowdhury, I. T. S. Li, T. T. M. Ngo, B. J. Leslie, B. C. Kim, J. E. Sokoloski, E. Weiland, X. Wang, Y. R. Chemla, T. M. Lohman, T. Ha, Defining Single Molecular Forces Required for Notch Activation Using Nano Yoyo. Nano Lett. 16, 1–20 (2016).

23. D. Seo, K. M. Southard, J. W. Kim, H. J. Lee, J. Farlow, J. U. Lee, D. B. Litt, T. Haas, A. P. Alivisatos, J. Cheon, Z. J. Gartner, Y. W. Jun, A mechanogenetic toolkit for interrogating cell signaling in space and time. Cell. 165, 1507–1518 (2016).

24. L. Morsut, K. T. Roybal, X. Xiong, R. M. Gordley, S. M. Coyle, M. Thomson, W. A. Lim, Engineering Customized Cell Sensing and Response Behaviors Using Synthetic Notch Receptors. Cell. 164, 780–791 (2016).

25. T. Verdorfer, H. E. Gaub, Ligand Binding Stabilizes Cellulosomal Cohesins as Revealed by AFM-based Single-Molecule Force Spectroscopy. Sci. Rep. 8, 9634 (2018).

26. K. Tiyanont, T. E. Wales, M. Aste-Amezaga, J. C. Aster, J. R. Engen, S. C. Blacklow, Evidence for Increased Exposure of the Notch1 Metalloprotease Cleavage Site upon Conversion to an Activated Conformation. Structure. 19, 546–554 (2011).

27. Y. Wu, C. Cain-Hom, L. Choy, T. J. Hagenbeek, G. P. de Leon, Y. Chen, D. Finkle, R. Venook, X. Wu, J. Ridgway, D. Schahin-Reed, G. J. Dow, A. Shelton, S. Stawicki, R. J. Watts, J. Zhang, R. Choy, P. Howard, L. Kadyk, M. Yan, J. Zha, C. A. Callahan, S. G. Hymowitz, C. W. Siebel, Therapeutic antibody targeting of individual Notch receptors. Nature. 464, 1052–1057 (2010).

28. M. Aste-Amézaga, N. Zhang, J. E. Lineberger, B. A. Arnold, T. J. Toner, M. Gu, L. Huang, S. Vitelli, K. T. Vo, P. Haytko, J. Z. Zhao, F. Baleydier, S. L’Heureux, H. Wang, W. R. Gordon, E. Thoryk, M. B. Andrawes, K. Tiyanont, K. Stegmaier, G. Roti, K. N. Ross, L. L. Franlin, H. Wang, F. Wang, M. Chastain, A. J. Bett, L. P. Audoly, J. C. Aster, S. C. Blacklow, H. E. Huber, Characterization of notch1 antibodies that inhibit signaling of both normal and mutated notch1 receptors. PLoS One. 5 (2010), doi:10.1371/journal.pone.0009094.

29. M. E. Fortini, D. Bilder, Endocytic regulation of Notch signaling. Curr. Opin. Genet. Dev. 19, 323–328 (2009).

30. B. Varnum-Finney, L. Wu, M. Yu, C. Brashem-Stein, S. Staats, D. Flowers, J. D. Griffin, I. D. Bernstein, Immobilization of Notch ligand, Delta-1, is required for induction of notch signaling. J. Cell Sci. 113 Pt 23, 4313–4318 (2000).

31. M. De Odrowąz Piramowicz, P. Czuba, M. Targosz, K. Burda, M. Szymoński, Dynamic force measurements of avidin-biotin and streptavdin-biotin interactions using AFM. Acta Biochim. Pol. 53, 93–100 (2006).

32. J. W. Weisel, H. Shuman, R. I. Litvinov, Protein-protein unbinding induced by force: Single-molecule studies. Curr. Opin. Struct. Biol. 13, 227–235 (2003).

33. R. Falk, A. Falk, M. R. Dyson, A. N. Melidoni, K. Parthiban, J. L. Young, W. Roake, J. McCafferty, Generation of anti-Notch antibodies and their application in blocking Notch signalling in neural stem cells. Methods. 58, 69–78 (2012).

34. A. Chopra, M. L. Kutys, K. Zhang, W. J. Polacheck, C. C. Sheng, R. J. Luu, J. Eyckmans, J. T. Hinson, J. G. Seidman, C. E. Seidman, C. S. Chen, Force Generation via β-Cardiac Myosin, Titin, and α-Actinin Drives Cardiac Sarcomere Assembly from Cell-Matrix Adhesions. Dev. Cell. 44, 87-96.e5 (2018).

35. M. S. Sakar, J. Eyckmans, R. Pieters, D. Eberli, B. J. Nelson, C. S. Chen, Cellular forces and matrix assembly coordinate fibrous tissue repair. Nat. Commun. 7, 11036 (2016).

36. A. M. Kabadi, P. I. Thakore, C. M. Vockley, D. G. Ousterout, T. M. Gibson, F. Guilak, T. E. Reddy, C. A. Gersbach, Enhanced MyoD-Induced Transdifferentiation to a Myogenic Lineage by Fusion to a Potent Transactivation Domain. ACS Synth. Biol. 4, 689–699 (2015).

37. H. E. Balcioglu, H. van Hoorn, D. M. Donato, T. Schmidt, E. H. J. Danen, The integrin expression profile modulates orientation and dynamics of force transmission at cell-matrix adhesions. J. Cell Sci. 128, 1316–1326 (2015).

38. K. Austen, P. Ringer, A. Mehlich, A. Chrostek-Grashoff, C. Kluger, C. Klingner, B. Sabass, R. Zent, M. Rief, C. Grashoff, Extracellular rigidity sensing by talin isoform-specific mechanical linkages. Nat. Cell Biol. 17, 1597–1606 (2015).

39. H. Li, M. Carrion-Vazquez, A. F. Oberhauser, P. E. Marszalek, J. M. Fernandez, Point mutations alter the mechanical stability of immunoglobulin modules. Nat. Struct. Biol. 7, 1117–1120 (2000).

40. W. J. Polacheck, C. S. Chen, Measuring cell-generated forces: A guide to the available tools. Nat. Methods. 13, 415–423 (2016).

41. Z. Yang, Z. Yu, Y. Cai, R. Du, L. Cai, Engineering of an enhanced synthetic Notch receptor by reducing ligand-independent activation. Commun. Biol. 3, 116 (2020).

42. W. R. Gordon, M. Roy, D. Vardar-Ulu, M. Garfinkel, M. R. Mansour, J. C. Aster, S. C. Blacklow, Structure of the Notch1-negative regulatory region: Implications for normal activation and pathogenic signaling in T-ALL. Blood. 113, 4381–4390 (2009).

43. M. D. Rand, L. M. Grimm, S. Artavanis-Tsakonas, V. Patriub, S. C. Blacklow, J. Sklar, J. C. Aster, Calcium depletion dissociates and activates heterodimeric notch receptors. Mol. Cell. Biol. 20, 1825–35 (2000).

44. J. E. Frith, G. D. Kusuma, J. Carthew, F. Li, N. Cloonan, G. A. Gomez, J. J. Cooper-White, Mechanically-sensitive miRNAs bias human mesenchymal stem cell fate via mTOR signalling. Nat. Commun. 9, 257 (2018).

45. S. Mangan, U. Alon, Structure and function of the feed-forward loop network motif. Proc. Natl. Acad. Sci. 100, 21 (2003)

46. D. Greber, M. Fussenegger, An engineered mammalian band-pass network. Nucleic Acids Res. 38 (2010), doi:10.1093/nar/gkq671.

47. P.C. Fridy, Y. Li, S. Keegan, M.K. Thompson, I. Nudelman, J.F. Scheid, M. Oeffinger, M.C. Nussenzweig, D. Fenyö, B.T. Chait, M.P. Rout, A robust pipeline for rapid production of versatile nanobody repertoires. Nature Methods, 11, 12 (2014)

48. P.D. Langridge, G Struhl, Epsin-dependent ligand endocytosis activates Notch by force. Cell, 171, 6 (2017).

49. B. Guo, BJ McMillan, SC Blacklow, Structure and function of the Mind bomb E3 ligase in the context of Notch signal transduction. Current Opinion in Structural Biology, 41, 38–45 (2016)

50. M. Messa, R. Fernández-Busnadiego, E. W. Sun, H. Chen, H. Czapla, K. Wrasman, Y. Wu, G. Ko, T. Ross, B. Wendland, P. De Camilli, Epsin deficiency impairs endocytosis by stalling the actin-dependent invagination of endocytic clathrin-coated pits. Elife, 3 (2014)

51. M. Kaksonen, A. Roux, Mechanisms of clathrin-mediated endocytosis. Nature Reviews Molecular Cell Biology, 19, 5 (2018)

52. D. Serwas, M. Akamatsu, A. Moayed, K. Vegesna, R. Vasan, J. M. Hill, J. Schöneberg, K. M. Davies, P. Rangamani, D. G. Drubin, Mechanistic insights into actin force generation during vesicle formation from cryo-electron tomography. Developmental Cell, 57, 9 (2022)

53. E. Seib, T. Klein, The role of ligand endocytosis in notch signalling. Biology of the Cell, 113, 10 (2021).

54. B. J. McMillan, B. Schnute, N. Ohlenhard, B. Zimmerman, L. Miles, N. Beglova, T. Klein, S. C. Blacklow. A tail of two sites: a bipartite mechanism for recognition of notch ligands by mind bomb E3 ligases. Molecular Cell, 57, 5 (2015)

55. S. Dengl, E. Hoffmann, M. Grote, C. Wagner, O. Mundigl, G. Georges, I. Thorey, K. G. Stubenrauch, A. Bujotzek, H. P. Josel, S. Dziadek. Hapten-directed spontaneous disulfide shuffling: a universal technology for site-directed covalent coupling of payloads to antibodies. The FASEB Journal, 29, 5 (2015)

56. J. B. McMahan, J. T. Ngo. A Genetically Encodable and Chemically Disruptable System for Synthetic Post-Translational Modification Dependent Signaling. bioRxiv 2022.05.29.493928; doi: https://doi.org/10.1101/2022.05.29.493928

57. F. Schwesinger, R. Ros, T. Strunz, D. Anselmetti, H. J. Güntherodt, A. Honegger, L. Jermutus, L. Tiefenauer, A. Plückthun, Unbinding forces of single antibody-antigen complexes correlate with their thermal dissociation rates. Proc. Nat’l. Acad. of Sci. U. S. A., 97, 18 (2000)

58. M. Trylinski, F. Schweisguth. Activation of Arp2/3 by WASp is essential for the endocytosis of Delta only during cytokinesis in Drosophila. Cell Reports, 28, 1, (2019)

59. J. D. Bloom, S. T. Labthavikul, C. R. Otey, F. H. Arnold, Protein stability promotes evolvability. Proc. Nat’l. Acad. of Sci. U. S. A. 103, 15 (2006)

60. L. Meloty-Kapella, B. Shergill, J. Kuon, E. Botvinick, G. Weinmaster. Notch ligand endocytosis generates mechanical pulling force dependent on dynamin, epsins, and actin. Developmental Cell, 12, 22 (2012)

