## Supplementary Information for v2 for "Tension-Tuned SynNotch Receptors for Synthetic Mechanotranduction and Intercellular Force Detection"

### MATERIALS AND METHODS

#### DNA constructs

DNA constructs were generated via standard cloning procedures, typically using Gibson assembly reactions. Inserts were generated by PCR amplification or acquired as custom synthesized DNA fragments (Integrated DNA Technologies, IDT). Plasmid backbones were linearized by digestion with restriction enzymes, or via amplification by PCR followed by template elimination using DpnI. Cloning of lentiviral backbones and sequences containing repeat regions was performed using NEB Stable Competent *E. coli* cells as the transformation host (C3040H, New England Biolabs). Plasmid DNAs and full sequence details for selected constructs have been deposited with AddGene (see below).

#### Mammalian cell culture

All mammalian cell lines were cultured in a humidified incubator maintained at 37 °C with 5% $CO_2$. Cell media were based on Dulbecco's Modified Eagle's Medium (DMEM), pH 7.0 - 7.4, containing high glucose and sodium pyruvate and without L-glutamine (SH30285.01, Cytiva, or similar), used with supplements as indicated below. Fetal bovine serum (FBS) was typically Cytiva Characterized Fetal Bovine Serum, Canadian Origin (SH30396.03, Cytiva), or Corning Regular Fetal Bovine Serum (35010CV, Corning). HEK293-FT cells (R70007, ThermoFisher) were cultured in DMEM with 10% FBS and supplemented with 1X nonessential amino acids (25025CI, Corning) and 1X GlutaMAX (3505006, ThermoFisher). HeLa cells (CCL-2, ATCC), C3H/10T1/2 cells (CCL-226, ATCC), and U-2 OS cells (92022711-1VL, Sigma-Aldrich) were cultured using the same medium but with added 1X penicillin-streptomycin solution (15140122, ThermoFisher).

#### DNA transfections

DNA transfections were carried out using Lipofectamine 3000 Reagent (L3000001, ThermoFisher) according to the manufacturer's instructions. For analyses involving transient transfection of HEK293-FT derived reporter cells (UAS:H2B-mCherry, described below), 25 ng of receptor-encoding plasmid was used in combination with 25 ng of a constitutively expressed fluorescent cotransfection marker per ~150,000 reporter cells. For transient expression of T2A-BFP fusions encoded on lentiviral backbones, 50 ng of plasmid without an additional cotransfection marker was used per ~150,000 reporter cells. For luciferase reporter measurements: 50 ng of plasmid DNA containing a UAS-regulated firefly luciferase (UAS-FLuc) reporter gene (AddGene # 46756) was used in combination with 1 ng of a plasmid encoding a constitutive NanoLuciferase gene as a cotransfection marker (pNL1.1.TK[*Nluc*/TK], Promega) per ~150,000 reporter cells.

For "low-pass" genetic circuits: three plasmids were cotransfected into cells, including 1) a miRNA "target" plasmid encoding a constitutively expressed mCerluean (mCer) gene tagged with an 8x-miRNA target sequence targeted by FF4 miRNA (AddGene plasmid # 26280, Nissim et al. *Mol Cell.* 54, (2014)), 2) a TRE-regulated mKate containing an intronic FF4 microRNA (miR-FF4), and 3) a receptor encoding plasmid in which TetR/tTA was used as the ICD. Cells were triply transfected with 25 ng of the mCer target plasmid, 25 ng of receptor-encoding plasmid, and 100 ng of the TRE-regulated mKate/miR-FF4 plasmid. Receptor activity was used to regulate TRE promoter activity and mKate/miR-FF4 for miRNA-mediated regulation of the targeted mCer gene.

For “band-pass” genetic circuits: individual constructs were designed to encode cross-swapped receptor-reporter pairings such that only cells containing both plasmids would exhibit receptor-mediated reporter gene activities; 25 ng of plasmid DNA encoding a Gal4-containing receptor and containing either a TRE-dsRed-Express2 gene cassette, or a TRE-Gal4-KRAB-T2A-dsRed-Express2 gene cassette, was used in combination with 25 ng plasmid encoding a tTA-containing receptor and containing a UAS-mCerulean reporter gene cassette.

Note that the utilized UAS-regulated FLuc sequence corresponds to a reporter plasmid originally described in 1994 and designated “5xGAL4-TATA-luciferase” (Sun *et al*. Genes Dev. 8, 21, (1994)). Reliable and reproducible signal detection was achieved using the plasmid. More recent commercial FLuc reporters with degron-fused enzymes and other enhancements reported to increase induction levels and response rates. These systems may be useful when increased sensitivity is required (see Promega Technical Manual #TM259: pGL4 Luciferase Reporter Vectors, Section 3).

**<u>Lentiviral production and transduction</u>**

For viral transductions, lentiviral particles were generated using a 2$^{nd}$-generation lentiviral vector system. HEK293-FT (ThermoFisher) cells were grown to approximately 90% confluence in a 6-well dish and transfected with 750 ng of construct-encoding transfer plasmid alongside 1.25 µg each of packaging (pPax2) and envelope encoding (pVSVG) plasmids using the Lipofectamine 3000 Reagent. Transfection media was replaced the following morning. Viral supernatants were harvested at 24 and 48 hour time points following media exchange. Supernatants were passed through low protein-binding 0.45 µm filters prior to immediate use, or storage by freezing at -80 °C. For viral transduction, cells were infected in growth media via addition of viral supernatant; viral media were replaced with fresh media 24 or 48 hours later.

**<u>Clonal reporter cell line generation</u>**

A reporter cell line containing a stably integrated Gal4-dependent reporter construct (UAS:H2B-mCherry) was generated by transfecting HEK293-FT cells with linearized reporter plasmid. The sequence used was based on pEV-UAS-H2B-Citrine (a gift from M. Elowitz, Caltech) following modification to H2B-mCherry-V5 fusion in place of H2B-citrine. The resulting plasmid is designated “pEV-UAS-H2B-mCherry.” The UAS promoter contains 14xGAL4 DNA binding sites upstream of the H2B-mCherry gene followed by a BGH poly-adenylation (pA) signal. The plasmid also contained a zeocin resistance marker expressed from the CMV promoter and terminated by a SV40-pA signal. The CMV promoter replaced the original SV40 promoter/origin to prevent large T-antigen mediated episomal replication in integrated cells. The UAS-H2B-mCherry plasmid was linearized by digestion outside of the expression cassettes and the resulting DNA was transfected into HEK293-FT cells. Cells were selected using 100 ug/mL zeocin at 72 hours post-transfection. Single clones were isolated by limited dilution into 96 well plates. An optimal reporter clone (designated "E5") for propagation on the basis of its low background H2B-mCherry levels and its high inducibility upon expression of a Gal4-based transcription factor (pcDNA3-Gal4-VP64). The clone maintained the geneticin (500 µg/mL) resistance of the parental HEK293-FT cells.

A U2OS reporter cell line containing a stable, virally-integrated UAS:dsRed-Express2 reporter was generated by transducing cells with viral constructs encoding UAS:dsRed-Express2 and PGK:PuroR gene cassettes. The construct were generated by modifying the LV-TRE-Empty-T2A-dsRedExpress2 vector (AddGene # 60623). Briefly, the TRE promoter was replaced with a UAS-based sequence containing 5xGAL4 binding sites and the original PGK:rTetR-IRES-PuroR cassette was replaced with a PGK:PuroR gene. The 5xUAS regulatory sequence from AddGene # 79123 was used. The gene cassettes did not contain terminating pA signals within its lentiviral backbone. Viral particles were produced as described above and stable integrants were selected using 0.5 µg/mL puromycin. Single clones were isolated and screened on the basis of low background and high inducibility, both as described above. The associated LV-UAS-DsRed-Express2-PGK-Puro plasmid, along with full sequence information, has been deposited to AddGene (see below).

**Sender and receiver cell line generation**

Stable receiver cell lines were generated via lentiviral integration of constructs EF1A-driven receptor genes fused to T2A-BFP at their C-termini and containing an IRES-driven hygromycin resistance marker. Gene fragments encoding receptors and T2A-BFP were ligated into the pLV-EF1A-IRES-Hygro vector backbone (AddGene # 85134). Receivers were generated by transducing HEK293-FT:UAS-H2B-mCherry reporter cells (Ngo Lab HEK reporter clone E5) or U2OS:UAS-T2A-dsRed-Express2 reporter cells (Ngo Lab U2OS reporter clone 1G4). Integrated clones were selected 48 hours after transduction using Hygromycin-B Gold (ant-hg-5, InvivoGen) at 75 ug/ml for HEK293-FT and at 200 ug/ml for U2OS cells. Transductions were performed in 24 well plates and cells were moved to 6 well plates for selection; initial antibiotic exposure was applied at no higher than ~25% cell confluence for efficient elimination of non-integrated cells. Selection with hygromycin typically required 10-12 days of growth in the presence of the drug, with media replenishment every 1-2 days. Transduced receiver cells were used as stably integrated pools without clone isolation.

For GFP-TMD-DLL1 and GFP-TMD-DLL1-Nbox senders, DNA fragments were cloned into a doxycycline-inducible "all-in-one" lentiviral backbone based on AddGene plasmid # 60627. Intracellular tails sequences are based on rat DLL1. The mutated GFP-TMD-DLL1-Nbox ligand contains a triple alanine substitution within the intracellular "N-box" motif (I**<u>KNT</u>**NKK to I**<u>AAA</u>**NKK). Ligand-encoding DNAs were inserted downtreams of the TRE promoter in place of the existing p65-MyoD-T2A-dsRed-Express2 fusion (AddGene # 60627). The vector backbone contained a PGK-regulated TetR followed by an IRES-drive puromycin marker for selection of integrated cells. Transduced HEK293-FT cells were selected 48 hours after transduction using puromycin at 0.5 ug/mL. Puromycin selection typically required 5-7 days, with regular replenishment of the antibiotic containing media. Transduced senders were used as stably integrated pools without clone isolation. HEK293-FT sender cells contained an integrated EF1A-anti-biotin-SNAP-TMD-DLL1-IRES-blasticidin construct are reported elsewhere (McMahan & Ngo 2022: doi:10.1101/2022.05.29.493928).

For myogenic differentiation: inducible C3H/10T1/2 cells were generated using a previously reported construct encoding a TRE-regulated p65-MyoD fusion (AddGene plasmid # 60627) (Kabadi *et al.* ACS Synth Biol. (2014)). Cells were transduced with lentiviral particles and selected using puromycin at 2 μg/ml 48 hours post-transduction. Single clones were isolated from drug selected pools via limited dilution into 96-well plates. Note that the referenced TRE-p65-MyoD construct also contains a PGK-driven cassette encoding the doxycycline-sensitive transcription factor rTetR. Thus, the generation, selection, clone isolation, and maintenance of the transduced C3H/10T1/2 cells were carried out using media containing tetracycline-free FBS (SH30070.03T from GE Healthcare, or 631106 from Takara, with similar results). Differentiation experiments using TGTs were also carried out under tetracycline-free conditions. Clones were tested using doxycycline (1 ug/mL) to verify TRE-dependent myogenic conversion.

**<u>Tension gauge tether (TGT) synthesis</u>**

To test the molecular tension requirements for NRR- and sNRR-based SynNotch activation, we immobilized SynNotch ligands to a tissue culture surface using tension gauge tethers (TGTs) using fabrication protocols described previously by the Ha lab. The method described in Wang et al., “Constructing modular and universal single molecule tension sensor using protein G to study mechano-sensitive receptors.” (*Sci Rep* 6, 21584 (2016); doi: 10.1038/srep21584) was followed closely, with modifications.

Custom single stranded DNA (ssDNA) sequences were ordered from Integrated DNA Technologies (IDT). For TGT experiments, receptors containing the E2 anti-fluorescein scFv were utilized in combination with the following FITC-containing oligonucleotide “ligand strand”:

Ligand strand: 5’-GGC CCG CAG CGA CCA CCC/**36-FAM**/-3’

The sequence above was paired with complementary ssDNAs as “tethering strands.” These strands contained amine functional groups located at varying positions along the ssDNA sequence; the amines were later modified with amine-reactive biotin as an immobilization handle (see below).

Amine containing complementary strands were ordered from IDT as follows:

For 12 pN TGTs: 5'-/**5AmMC6**/GGG TGG TCG CTG CGG GCC-3'
For 43 pN TGTs: 5'-GGG TGG TCG C/**iAmMC6T**/G CGG GCC-3'
For 54 pN TGTs: 5'-GGG TGG TCG CTG CGG GCC/**3AmMO**/-3'

Modified positions are shown in bold in the sequences, as designated based on their IDT modification codes. Further details regarding these modifications are provided below.

| *IDT code* | *Name* | *Structure* |
| --- | --- | --- |
| 36-FAM | 3' 6-FAM (6-fluorescein) | 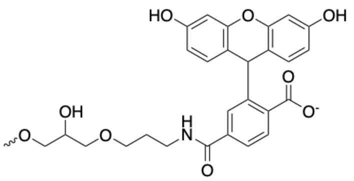 |
| 5AmMC6 | 5' Amino Modifier C6 | 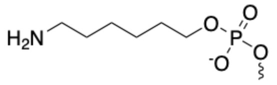 |
| iAmMC6T | Int Amino Modifier C6 dT | 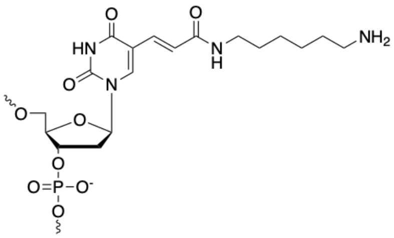 |
| 3AmMO | 3' Amino Modifier | 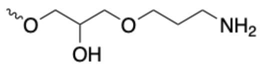 |

Biotin-modified tethering strands were generated by reacting amine groups with an NHS-ester functionalized $PEG_{12}$-biotin (EZ-Link NHS-$PEG_{12}$-biotin; 21312, ThermoFisher). To biotinylate the amine-modified tethering strands we followed a previously reported step-by-step protocol published by Wang and coworkers (*Scientific Reports*, v. 6, 21584 (2016); see "Materials and Methods: Preparation of complementary ssDNA-biotin."

Batch preparations of biotinylated tethering strands were stored at 100 uM concentration in a 10 mM Tris buffer (pH 7.4) containing 150 mM NaCl. Dried FITC-conjugated ligand strand received from the supplier was dissolved to 100 uM concentration in a buffer of the same composition. Aliquots the ssDNA solutions were stored at -20 °C until use. To prepare dsDNA TGT duplexes, FITC-conjugated ligand strand DNA was combined with individual tethering strand sequences at 1:1 (v/v) mixtures. Annealing reactions were carried out by overnight incubation at 4 °C resulting in 50 uM dsDNA TGT stock solutions.

**Tension gauge tether (TGT) assay**

Preparation of cells for flow cytometry analyses was typically done using plastic, non-tissue culture treated 96-well microwell plates. When preparing cells for imaging TGT-induced responses, plates with non-coated coverslip glass bottoms were used. Coating, immobilization, and wash solutions were applied to microwell plates by multichannel pipetting, typically using 100 uL volumes.

To immobilize TGTs, microwells surfaces were treated with solutions containing a mixture of 10 µg/ml biotinylated BSA (A8549, Sigma-Aldrich) in combination with 10 µg/ml fibronectin (AAJ62380LB0, Fisher Scientific) in PBS for 1 hour at room temperature prior to washing three times with PBS. Neutravidin (31000, ThermoFisher) was then immobilized to the biotin-coated surfaces by treating with a solution diluted to 100 µg/ml in PBS. Binding proceeded for 30 min at room temperature prior to washing three times with PBS. TGT from the prepared 50 uM dsDNA stocks were then diluted to 100 nM in PBS and added to the neutravidin-treated microwells. TGT binding to neurtravidin surfaces was allowed to proceed for 1 hour at room temperature prior to washing with PBS three times.

For "upper limit" $T_{tol}$ controls, fluorescein conjugated to biotin via a short PEG linker was used (22030, ThermoFisher). In this case, the "upper limit" $T_{tol}$ is defined as the tension level required to disengage the biotin-streptavidin bonds (generally estimated as 100 pN or more: Lee et al. *Micron*. 38 (2007), Johnson and Thomas, *Biophys. J.* 114, 9 (2018)). Attached to neutravidin coated cells was done using 100 nM solution in PBS. Binding proceeded for 30 min room temperature prior to washing with PBS three times.

Cells prepared for TGT analysis were transfected in suspension mixtures as follows: HEK293-FT reporter cells (UAS:H2B-mCherry) cells were trypsinized and the quenching media was removed prior to resuspending cells in fresh media. Prepared DNA-lipid transfection mixtures were then added to cell suspensions and gently mixed seeding with TGT-containing wells using ~30,000 cells per well. Cells were analyzed for receptor-mediated reporter expression 48 hours later. Note cell differentiation experiments were done using transduced C3H/10T1/2; cells were analyzed for myogenic conversion at 72 hours following TGT exposure, as indicated in the section describing the myogenic differentiation assay.

**Coculture *trans*-activation assays**

Non-treated tissue culture 96-well plates were coated using 10 ug/ml fibronectin (FN) solutions in PBS for 1 hour at room temperature. Wells were then washed three times with PBS and allowed to dry. Sender and receiver cell populations were then trypsinized using a pre-warmed 0.25% trypsin solution without EDTA. Cultures were quenched with warm media prior to centrifugation, aspiration, and resuspension in fresh warm media. Cell densities were individually measured using an automated cell counter (Countess II, ThermoFisher) and suspensions were inspected for homogenous cell distributions to ensure minimal cell clumping. Sender and receiver cells were combined at 2:1 ratios (sender:receiver) in FN-coated 96 well microwells and cultivated for 24 hours before analysis. The following cell numbers were used for cocultures at the 96 well scale: 20,000 HEK293-FT receivers with 40,000 HEK293-FT senders, or 10,000 U2OS receivers with 20,000 HEK293-FT senders.

HEK293-FT GFP-DLL1/Nbox ligand cells were preincubated with 100 ug/ml doxycycline for 24 hours prior to coculturing with receiver cells, during which doxycycline was maintained in the coculture medium at the same concentration. For dynamin inhibition, cells were exchanged into pre-warmed Gibco Opti-MEM I Reduced Serum Medium (31985062, ThermoFisher) containing 80 uM dynasore (D7693-5MG, Sigma-Aldrich) and 2 nM biotin-FITC at approximately 12 hours post-cell seeding.

**Imaging and preparation of microwell patterned *trans*-activations**

For microwell-patterned *trans*-activations we used non-treated 35 mm glass-bottom imaging dishes (P35G-1.0-20-C, MatTek). Dishes coated with fibronectin using 20 ug/ml fibronectin solutions in PBS with overnight incubation at 37 °C. Surfaces were washed three times with PBS and allowed to dry at 4 °C. A 3-well silicone insert containing two defined cell-free gaps (80369, ibidi) was applied to the center of the coated dishes using sterile forceps. Gentle pressure was applied to ensure attachment of the insert with the glass via its adhesive coating. Cells were then seeded into chamber wells at ~35,000 cells per well suspended in roughly 70 uL culture medium (corresponding to cell densities of ~500k/mL). Care was taken to ensure even cell disbursement in each well (media and solutions were pre-warmed to 37 °C and cells were mixed via gentle pipetting). After seeding cells, 1.5 mL of culture medium was added to the area surrounding the insert to limit evaporation from well interiors. Dishes were capped and cells were returned to the incubator for attachment and growth. When confluence within the chambers reached ~90% (~ 24 hours later), microwell inserts were carefully removed using sterile forceps. Following an additional ~18 hours of growth, media were exchanged with fresh media containing 2 nM biotin-FITC. After ~24 hours, cells were processed for imaging as indicated below.

Reporter (UAS-H2B-mCherry) expression levels were detected at sender-receiver interfaces and recorded using epifluorescence microscopy. Prior to imaging, senders in the patterned cocultures were fluorescently marked by labeling with a cell impermeant SNAP tag dye (SNAP-Surface AlexaFluor647, S9136S, New England Biolabs). The dye was diluted to 4 μM concentration in pre-warmed culture media (1:250 dilution from a 1 mM stock in DMSO) and the solution was vortexed to ensure full dye dissolution. The mixture was incubated at 37 °C for an additional 15 minutes before applying to cells. Labeling was initiated by adding ~600 uL of dye solution per 35 mm imaging dish followed by incubation at 37 °C for 30 min. Cells were rinsed with warm media three times prior to an additional 30 minutes in the incubator in fresh dye-free media. For imaging, cells were exchanged into FluoroBrite DMEM imaging media (A1896701, ThermoFisher) containing supplements similar to growth media as described above. Hoechst 33342 was used as a nuclear counterstain at 1 uM concentration.

**Protein expression and purification**

For expression of soluble anti-NRR scFv-Fc fusion: DNA encoding an scFv corresponding to anti-Notch1-NRR originally described by Wu and coworkers (Wu et al. *Nature*, 464, (2010)) was subcloned into the pBIOCAM5 vector backbone (Addgene # 39344). The fragment was inserted between *NcoI* and *NotI* restriction sites following removal of the existing insert. The resulting plasmid encodes an in-frame secreted fusion construct containing the scFv followed by a human Fc region and C-terminal 6xHis and 3xFLAG tags. Transient expression of this construct was carried out by transfection of HEK293-FT in 100 mm tissue culture dishes. The secreted protein was collected by harvesting conditioned media over the course of seven days. The collected media was stored at 4 °C and combined prior to passage through a low protein binding 0.2 µm filter. The filtrate pH was adjusted by adding pH 8 Tris-buffered saline solution to a final concentration of Tris at 20 mM and 300 mM NaCl. The scFv-Fc-6xHis-3xFLAG fusion was then purified using the Ni-NTA Fast Start Kit (30600, Qiagen) following manufacturer's protocol. Imidazole was removed from protein eluents by dialysis against PBS at 4 °C using 10,000 MWCO Slide-A-Lyzer Dialysis Cassettes (66380, ThermoFisher). Purity was assessed by staining SDS-PAGE gels with GelCode Blue Stain Reagent (24590, ThermoFisher) with colorimetric quantification against a BSA standard serially diluted from a 2 mg/mL stock solution (23209, ThermoFisher).

For expression of eGFP: DH10B *E. coli* (EC0113, ThermoFisher) carrying plasmid DNA encoding an arabinose-inducible His6x-eGFP (EGFP-pBAD, AddGene # 54762) were selected on LB-agar containing 100 ug/ml ampicillin. A single clone was transferred to 5 mL LB media with ampicillin and grown overnight at 37 °C with agitation. The following day, 4 mL of culture was used to inoculate 200 mL of LB-ampicillin. The culture was allowed to grow until OD = ~0.7, at which point the flask was transferred to a 30 °C incubator for 30 min prior to adding arabinose to a final concentration of 0.2% (w/v). Protein expression proceeded at 30 °C with shaking at 250 rpm for 4 hours. Cells were harvested by centrifugation and stored as frozen pellets at -20 °C until purification. His6x-eGFP was purified using the Ni-NTA Fast Start Kit (30600, Qiagen) according to the manufacturer's protocol. Imidazole was removed from protein eluents by gel filtration with PD-10 Sephadex G-25 resin columns (GE17-0851-01, Millipore-Sigma) followed by dialysis against PBS using 10,000 MWCO Slide-A-Lyzer Dialysis Cassettes (66380, ThermoFisher). The EGFP-pBAD vector was maintained in cells using 100 µg/ml ampicillin. The purity of the isolated protein was assessed by staining SDS PAGE gels using GelCode Blue Stain Reagent; GFP concentration was determined using the bicinchoninic acid protein assay (BCA Protein Assay Kit, 23225, ThermoFisher).

For expression of mono-biotinylated GFP: EGFP-pBAD plasmid was modified by inserting an in-frame DNA segment encoding a tandem linked AviTag-ALFA tag

(GLNDIFEAQKIEWHE-GSGGS-RLEEELRRRLTE-GSGTKAS) in between His6x and eGFP (pBAD-6xHis-AviTag-ALFA-eGFP). Expression was carried out as described for eGFP with modifications. An *E. coli* strain with inducible biotin ligase (BirA) was used as the expression host (AVB101, Avidity). At the time of arabinose induction, d-biotin was added to 50 uM along with isopropyl β-d-1-thiogalactopyranoside (IPTG) at 1.5 mM (to induce BirA overexpression in AVB101). AVB101 was maintained using 10 μg/ml chloramphenicol.

**Antibodies and labeling proteins**

Commercial antibodies and labeling proteins were used at the indicated dilutions:

| *Antibody or Labeling Protein* | *Dilution* | *Supplier* |
|---|---|---|
| Mouse anti-myc-AlexaFluor647 | 1:200 | sc-40 AF647, Santa Cruz Biotechnology |
| Mouse anti-myc-Alexa Fluor488 | 1:200 | MA1-980-AF488, ThermoFisher |
| Mouse anti-myc | 1:200 | AHO0062, ThermoFisher |
| Mouse anti-myosin heavy-chain | 1:100 | MAB4470, R&D Systems |
| Mouse anti-NRR E6 | 1:1,000 | Ab00175-1.1, Absolute Antibody (*51*) |
| Mouse anti-Notch1 E4 (C-term) | 1:500 | sc-373944, Santa Cruz Biotechnology |
| Rabbit anti-RFP | 1:1000 | 600-401-379, Rockland Immunochem. |
| GFP-Booster AlexaFluor647 | 1:500 | gb2AF647-50, ChromoTek/Proteintech |
| Mouse anti-GAPDH | 1:3,000 | MA5-15738, ThermoFisher |
| Direct-Blot HRP anti-GAPDH | 1:4,000 | 607904, BioLegend |
| Anti-mouse-HRP | 1:3,000 | 7076, Cell Signaling Technology |
| Anti-rabbit-HRP | 1:3,000 | 1721019, Bio-Rad |
| Goat anti-mouse-AlexaFluor488 | 1:2,000 | A-11001, ThermoFisher |
| Rabbit anti-mouse-AlexaFluor647 | 1:1,000 | A-21239, ThermoFisher |
| Goat anti-human-AlexaFluor647 | 1:1,000 | A-21445, ThermoFisher |
| Dll4Fc (human Fc) | 5 μg/mL | A42511, ThermoFisher |
| WGA-AlexaFluor647 | 4 μg/mL | W32466, ThermoFisher |

In-house purified proteins were prepared as described in the preceding section and used at the concentrations indicated below:

| *Purified Protein* | *Concentration* |
|---|---|
| Anti-Notch1-NRR-scFv-Fc | 1 μg/mL for cell labeling solutions |
| eGFP | 20 μg/mL for cell labeling solutions |
| biotinylated-eGFP | 100 nM for ligand immobilization solutions |

**Immunofluorescence staining of fixed cells**

Cells grown in medium were rinsed with PBS prior to fixation with a pre-warmed formaldehyde solution diluted to 4% (v/v) in PBS from a vial containing 16% (v/v) stock solution (28906, ThermoFisher). Fixation was allowed to proceed for 10 min at room temperature. Cells were then rinsed 3 times with PBS followed by blocking for 1 h at room temperature using a BSA solution at 5% (w/v) in PBS. Staining was carried out for

1 hr at room temperature using primary antibody or DLL4-Fc diluted in 1% (w/v) BSA solution in PBS at the amounts indicated above. Cells were rinsed three times using PBS for 5 minutes per rinse prior to staining with secondary antibodies at the dilutions indicated above. When necessary, cells were permeabilized with a PBS solution containing 0.2 % Triton X-100 (v/v).

For visualizing receptor recycling from the cell surface, live cells were stained with soluble purified GFP prior fixation. Staining was carried out for either 45 min at 4 C (to prevent receptor internalization, Fig. S1B), or under internalization-permissive conditions for 30 min at 37 C (Fig. S1C). Labeled cells were then washed 3 times using PBS for 5 minutes each, at either 4 C or room temperature, respectively. Cells were fixed using 4% formaldehyde prior to surface-selective labeling against the NRR. Immunofluorescence detection of the NRR was carried out using purified anti-Notch1-NRR-scFv-Fc in all cases except when co-labeling cells with Dll4-Fc, which also contains the human Fc region. Thus, Dll4-Fc was used in combination with mouse anti-NRR-E6 when co-labeling Notch1 expressing cells (Fig. S4E). Surface selective NRR detections were performed using fixed cell specimens without permeabilization, both as described above.

### Preparation of cell lysates and immunoblotting

Cells were prepared for immunoblot analysis by rinsing with PBS followed by direct lysis using a 1X LDS-PAGE loading buffer solution (NP0007, Invitrogen). Lysates were then sonicated or sheared (by passage through a thin gauge syringe needle) to reduce lysate viscosities prior to clarification by centrifugation. Subsequent immunoblot preparation, including gel electrophoresis and membrane transfer, were carried out using standard procedures. Nitrocellulose membranes were blocked using solutions containing 5% non-fat dry milk (w/v) or 5% BSA (w/v), both diluted in PBS-Tween 20 (PBS-T, 0.1%, v/v). Blocking solutions were selected based on the specifications provided by primary antibody vendors. Primary and secondary antibody probing solutions were prepared at the dilutions indicated above. Detection of labeled antigens was carried out by chemiluminescence using the SuperSignal West Pico PLUS Chemiluminescent Substrate (34580, ThermoScientific). Membranes were stripped using the Restore Western Blot Stripping Buffer (21059, ThermoFisher) and re-blocked prior to re-probing.

### Luciferase assay for surface-adhered ligand and EDTA treatment

HEK293FT cells were triply transfected with three constructs: a UAS-regulated firefly luciferase reporter plasmid (AddGene plasmid # 46756) , a receptor encoding plasmid, and a cotransfection marker based on a plasmid encoding a constitutively expressed NanoLuciferase gene (pNL1.1.TK[*Nluc*/TK], Promega).

To investigate ligand-mediated activation of NRR- and sNRR-based anti-FITC SynNotch receptors, transfected cells were grown in fibronectin (FN)-coated microwells with or without immobilized biotin-FITC. To coat surfaces, non-treated tissue culture microwells were treated with protein-containing solutions as follows: for control surfaces without ligand, well were treated with PBS solutions containing FN at 10 µg/mL; for ligand-containing surfaces, wells co-treated using PBS solutions containing FN at 10 µg/mL in combination with 10 µg/ml biotinylated BSA. Coating proceeded for 1 hour at room temperature prior to rinsing with PBS. Wells containing biotinylated-BSA were subsequently treated with neutravidin followed by biotin-FITC, as described in the TGT methods.

To evaluate dependence on ADAM10 and gamma-secretase cleavage, cells were treated with either the broad-spectrum metalloprotease inhibitor batimastat (BB-94 SML-0041, Sigma-Aldrich) at 20 µM, or using the gamma secretase inhibitor Compound E (15579, Cayman Chemical) at 400 nM concentration. When using the gamma secretase DAPT (N-[N-(3,5-difluorophenacetyl)-L-alanyl]-S-phenylglycine t-butyl ester), 10 µM concentrations were applied (HY-13027, MedChemExpress). To evaluate the requirement of ligand-mediated tension, cells were treated with soluble fluorescein (fluorescein, sodium salt, 46960-25G-F, Sigma-Aldrich) in culture medium at the concentrations indicated within the captions. Luciferase expression levels were measured 24 hours following transfection using the Nano-Glo Dual Luciferase Reporter Assay System (N1620, Promega) according to the manufacturer’s protocol.

To investigate receptor activation dependent on calcium chelation, transfected cells were grown for 24 h before treatment with either PBS or with PBS containing 0.5 mM EDTA for 15 min at 37 °C (*29*). Note that PBS without added calcium and magnesium was used. Following EDTA treatment, the chelator was quenched by the addition of growth media to the cell suspensions. Cells were incubated under growth conditions for an additional 6 hours prior to luciferase measurement using the Nano-Glo Dual Luciferase Reporter Assay System as described above.

### Myogenic differentiation assay

Clonal C3H/10T1/2 reporter cells with stably integrated TRE:p65-MyoD-T2A-dsRed were transduced with lentiviral constructs encoding NRR- or sNRR-based receptors. At 48 hours post-infection, cells were treated with EDTA-free trypsin and plated on TGT- and FN-coated 8 well chambered glass coverslips (80826, ibidi) at 50,000 cells per well. Following 72 hours of growth on TGTs, cells were fixed and permeabilized and myogenic differentiation was assessed by immunofluorescence detection of myosin heavy-chain expression in combination with DAPI counterstaining for identification of multinucleated fibers.

### Flow cytometry

Cells were analyzed at roughly 48 hours after transfection using an Attune NxT flow cytometer (ThermoFisher). Live cells were identified by setting minimum FSC-A and SSC-A gating thresholds (Fig. S15A). Singlet cells were then identified by setting a polygon gate on FSC-A versus FSC-H (Fig. S15B). Positively transfected cells were identified via detection of their co-transfection or self-cleaving ‘T2A’ fluorescent protein markers, gated based on the 99$^{th}$ percentile of fluorescence signal under these conditions detected from non-transfected control cells (Fig. S15C). To improve detection of cells expressing both a receptor of interest and cotransfection marker, positively transfected cells were then gated for those bearing marker fluorescence emission at levels above the median of all transfected cells. In the case of virally transduced cells analyzed in Fig. 3, fluorescent marker gating was not applied to the high efficiency of lentiviral transduction for the associated analyses. Receptor cells in coculture assays were gated by BFP+. Experimental groups were analyzed for the fraction of cells representing positively activated reporter gene expression, defined based on comparison to control reporter cells (lacking any receptor) using emission intensities from the 99th percentile of control cells analyzed under identical excitation and detection parameters. Flow cytometry data were analyzed and quantified using R with the open source ggCyto software (https://github.com/RGLab/ggcyto). An example of the gating procedures utilized is provided below in **Fig. S15**.

### sNRR construct design and numbering convention for mutant domains

Mutations were introduced within the sNRR domain at positions residing within scFv or NRR regions. Amino acid positions within the scFv region were numbered order of insertion from 1 to 244. For the NRR, positions were identified based on their full-length Notch-1 numbering number designations. These conventions are depicted in the sNRR-SynNotch diagram below.

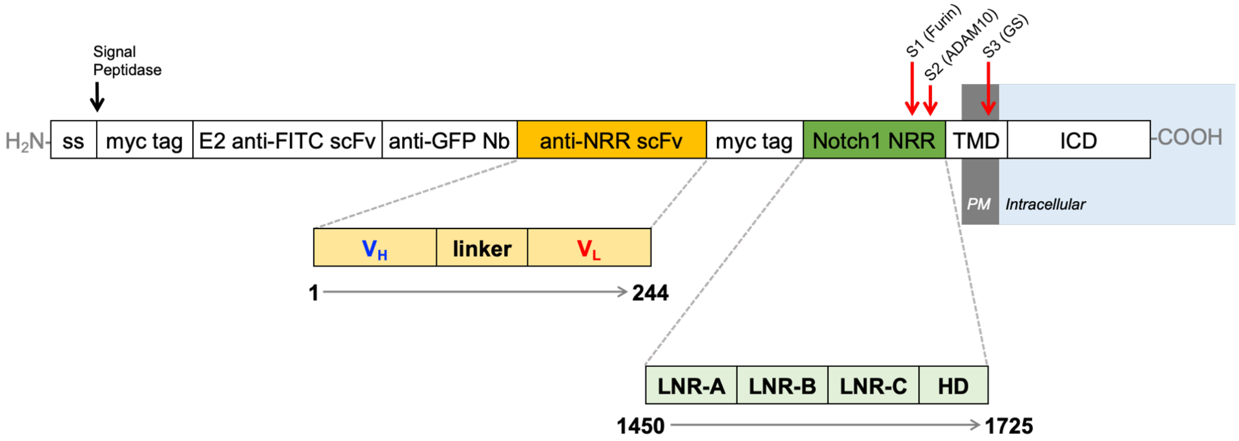


Note that the second myc tag between the anti-NRR scFv and NRR appears only in sNRR; NRR-SynNotch sequences contain only 1 myc tag, positioned N-terminally to the anti-FITC domain. Abbreviation and designations in the diagram above are defined as follows: **ss** - signal sequence peptide from CD8a, $\mathbf{V_H}$ - heavy chain variable region, $\mathbf{V_L}$ - light chain variable region, **Nb** - Nanbody (the LaG16 Nb was used for receptor *trans*-activation), **HD** - heterodimerization domain.

### Anti-Notch1 scFv sequence used in sNRR generation

The scFv sequence for the anti-Notch 1 NRR is provided below with the associated numbering used in the designation of mutated scFv positions. Blue letters designate $V_H$

residues, black indicates linker residues, red correspond to $V_L$ residues. Mutated positions are bolded and highlighted in yellow.

```
1        10        20        30        40        50
|        |         |         |         |         |
EVQLVESGGGLVQPGGSLRLSCAASGFTFSSYWIHWVRQAPGKGLEWVAR

51       60        70        80        90        100
|        |         |         |         |         |
INPPNRSNQYADSVKGRFTISADTSKNTAYLQMNSLRAEDTAVYYCARGS

101      110       120       130       140       150
|        |         |         |         |         |
GFRWVMDYWGQGTLVTVSSGGSSRSSSSGGGGSGGGGDIQMTQSPSSLSA

151      160       170       180       190       200
|        |         |         |         |         |
SVGDRVTITCRASQDVSTAVAWYQQKPGKAPKLLIYSASFLYSGVPSRFS

201      210       220       230       240
|        |         |         |         |
GSGSGTDFTLTISSLQPEDFATYYCQQFYTTPSTFGQGTKVEIK
```

**Sequence details for DNA constructs**

Plasmid DNA and full sequence information for the following constructs have been deposited to AddGene under the listed deposition numbers

| *Deposited Plasmids* | *AddGene #* | *Notes* |
|---|---|---|
| LV anti-FITC-LaG16-NRR-T2A-BFP | ###### | WT NRR |
| LV anti-FITC-LaG16-sNRR-T2A-BFP | ###### | WT sNRR |
| LV anti-FITC-LaG16-Y186F-T2A-BFP | ###### | |
| LV anti-FITC-LaG16-Y186A-T2A-BFP | ###### | |
| LV anti-FITC-LaG16-Y192A-T2A-BFP | ###### | |
| LV anti-FITC-LaG16-R103A-T2A-BFP | ###### | |
| LV anti-FITC-LaG16-Y186F/Y229A-T2A-BFP | ###### | 2x mutant sNRR |
| LV TRE-GFP-TMD-DLL1 | ###### | |
| LV TRE-GFP-TMD-DLL1-Nbox | ###### | |
| LV UAS-DsRed-Express2-PGK-Puro | ###### | UAS/Gal4 reporter |

*Note: AddGene deposition numbers are TBD as of 08/08/2022*

### Supplementary Figures

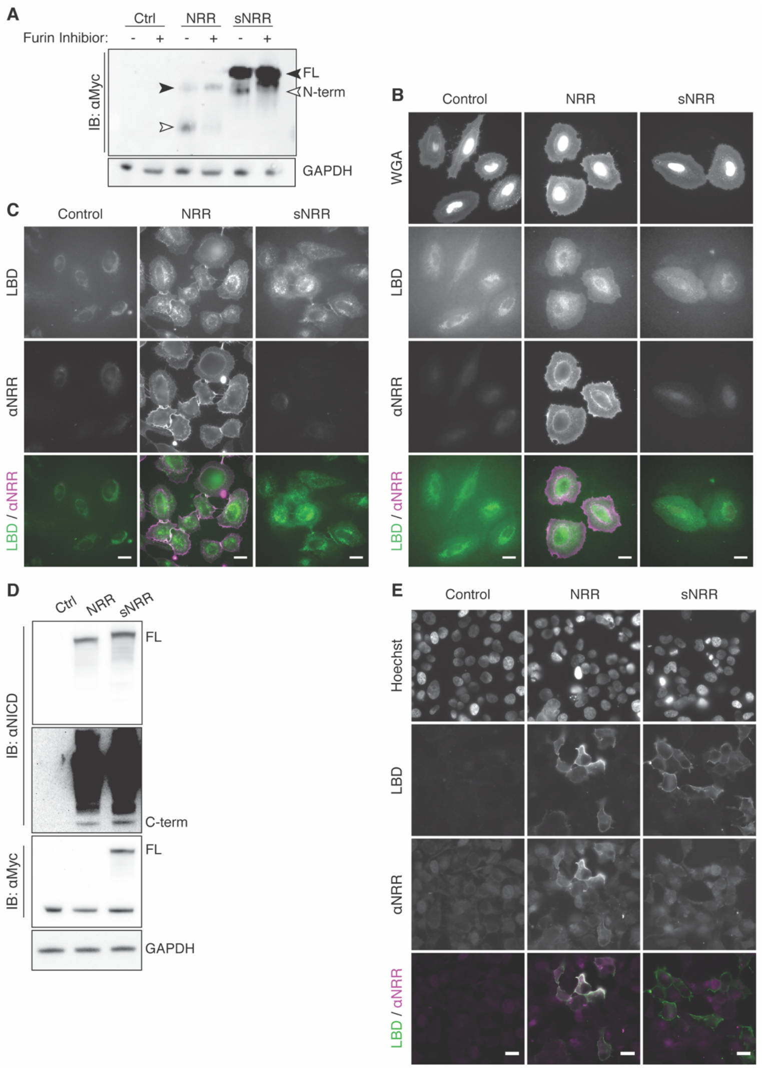


**Fig. S1. Trafficking and processing of receptors.** (**A**) Immunoblotting shows that NRR- and sNRR-based SynNotch receptors are processed into a noncovalent heterodimer by furin convertase. Furin inhibition induces the loss of cleaved N-terminal fragments (N-term, white arrowheads) and accumulation of full-length receptors (FL, black arrowhead). sNRR-based SynNotch receptors possess two myc tags N-terminal to the furin cleavage site, while NRR-based SynNotch receptors have one myc tag. (**B**) Immunostaining of SynNotch receptors shows surface expression and NRR accessibility in HeLa cells. Receptors express a GFP-binding nanobody as the LBD, which is stained using soluble GFP (green). Wheat germ agglutinin (WGA) stains cell membranes. Control cells are nontransfected. (**C**) Similar to (B), except GFP is used to stain the LBD at 37 °C, rather than 4 °C, to allow receptor internalization. Subsequent fixation and anti-NRR staining reveals that surface stained receptors are constitutively internalized and recycled from the cell surface, as evident by internalized LBD staining that does not colocalize with surface NRR staining. (**D**) Immunoblotting showed that NRR- and sNRR-based full-length recombinant human Notch1 receptors (rather than SynNotch receptors) are processed into noncovalent heterodimers, as evidenced by detection of the C-terminal fragment. Here, only sNRR receptors express a Myc epitope. (**E**) Similar to (B), in cells expressing full-length recombinant Notch1 receptors, rather than synthetic Notch receptors. Labeling the LBD with soluble Dll4/Fc shows receptors at the cell surface. Incorporation of a sNRR domain blocks reactivity to staining with exogenous anti-NRR .

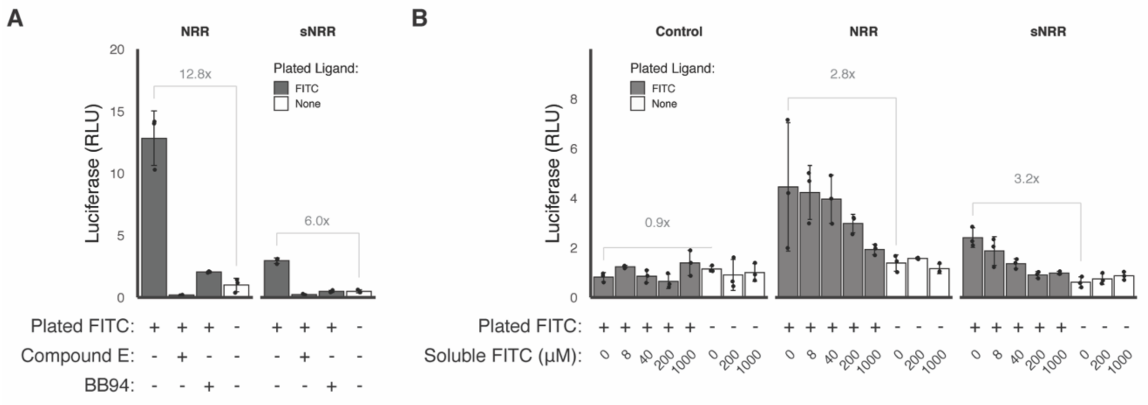


**Fig S2. Ligand-mediated activation of receptors.** (**A-B**) Cells expressing NRR- and sNRR-based receptors containing Gal4-VP64 ICDs were cultured on control surfaces containing fibronectin only, or grown in fibronectin- and FITC ligand-coated wells. Receptor activation induced the activation of a Gal4-dependent firefly luciferase reporter gene. (A) Plated ligand induced luciferase reporter activity, which was reduced upon treatment with BB94 (20 µM; for metalloproteinase inhibition) or with Compound E (400 nM; a gamma secretase inhibitor). (B) Treatment with varying amounts of soluble fluorescein resulted in competitive inhibition against surface ligands at high soluble fluorescein levels. Soluble ligands alone did not induce signaling upregulation; soluble ligands cannot offer tensile resistance to unravel the NRR or sNRR domain. Control cells express NRR-based SynNotch receptors with a tTA-based ICD, which is mismatched with respect to the implemented UAS-luciferase reporter gene.

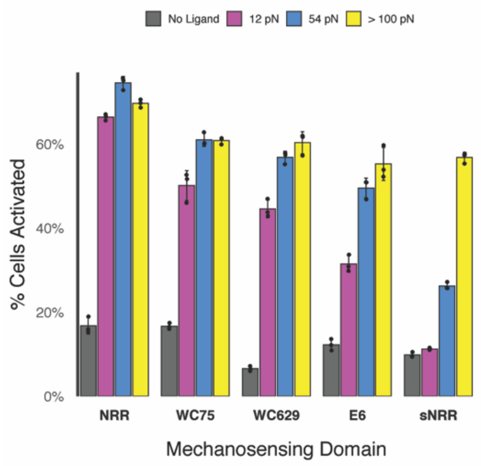


**Fig. S3. Additional NRR-scFv pairs tested on TGT's.** In addition to the original scFv tested in the sNRR domain (right set of bars; gray), three additional Notch1 NRR-binding antibodies were expressed as scFv receptor functions similar to sNRR. The tensile sensitivity of these receptors were evaluated using FITC-conjugated TGTs via transient expression in reporter HEK293-FT cells (UAS-H2B-mCherry). Quantified signaling responses are shown above; individual samples were analyzed using 50,000 cells or more.

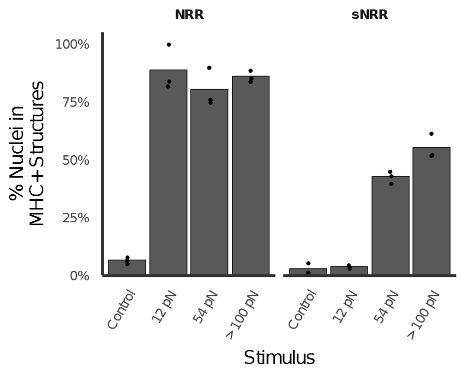


**Fig S4. Tension-induced myogenic differentiation.** C3H10T1/2 fibroblasts expressing NRR- or sNRR-based SynNotch receptors induce p65-MyoD expression upon activation which facilitates myogenic differentiation. Extents of myogenic differentiation were quantified as percentages of DAPI-labeled nuclei localized to cellular structures with positive-costaining against Myosin Heavy Chain (MHC). Data represent quantifications from 3 immunostained fluorescence micrographs per group using imagings containing 100 cells or more.

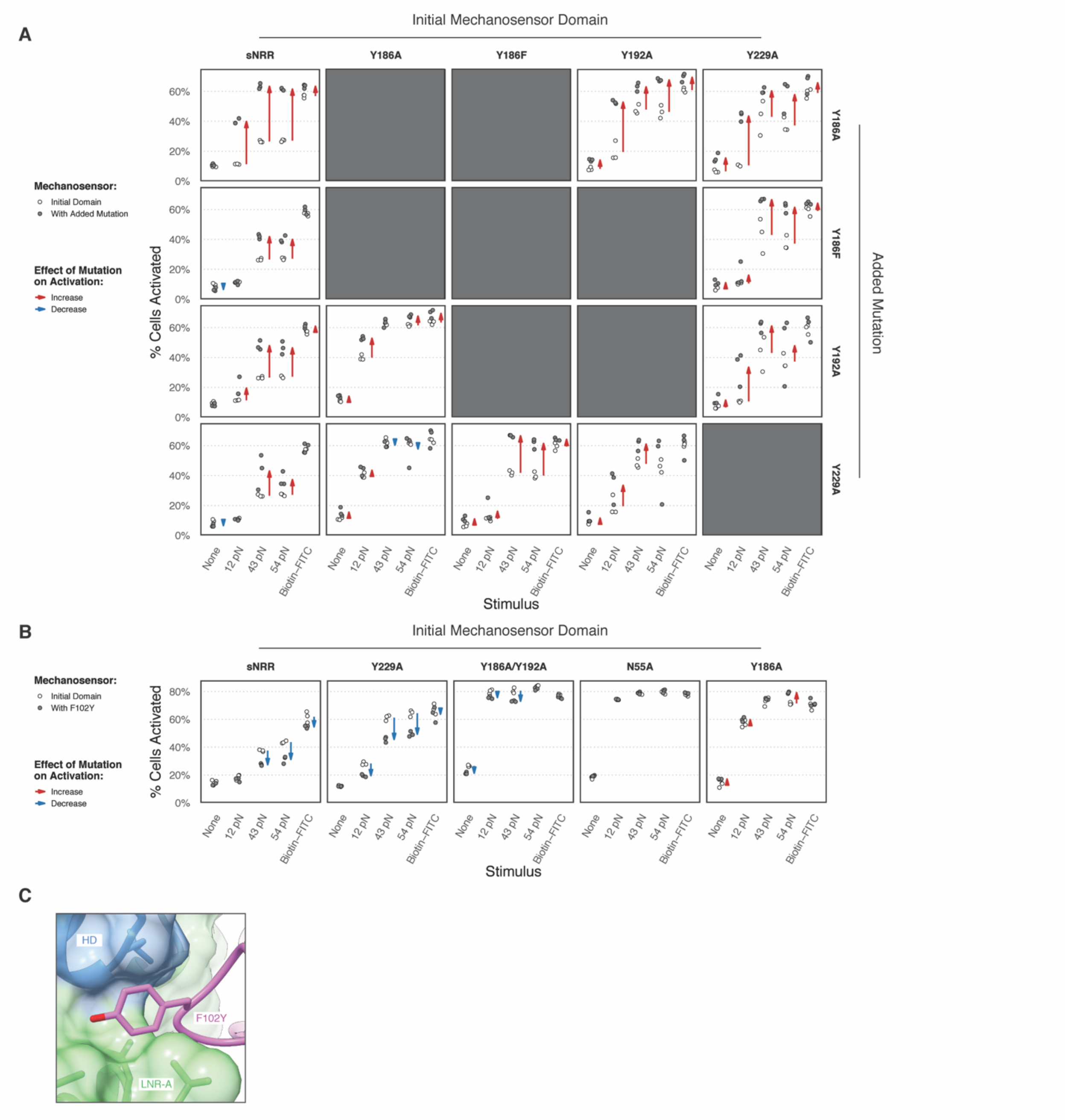


**Fig. S5. Tuning mechanical strength through double mutation or strengthening mutations.** (**A**) An extension of the data set presented in Figure 2D, detailing the effect of mutations on receptor strength. Each individual plot depicts the effect of an added mutation (gray circles on plots, vertical axis of grid) on an initial mechanosensing domain (open circles on plots, horizontal axis of grid). Arrows denote the change in mean activation upon mutation, for changes of magnitude > 5%. Mutations that weaken the receptor are expected to increase activation (red), while mutations that strengthen the receptor are expected to reduce activation (blue). (**B**) Effect of the F102Y mutation, represented as in (A). Addition of F102Y generally increases receptor strength, except when added to the Y186A mutation, which is immediately next to F98 in three-dimensional space.. (**C**) Modeled mutagenesis of F102Y. F102Y adds a hydroxyl group at the para-position. Rotamer modeling predicts this group would sterically fit at the binding interface and would be solvent-exposed (PDB 3L95).

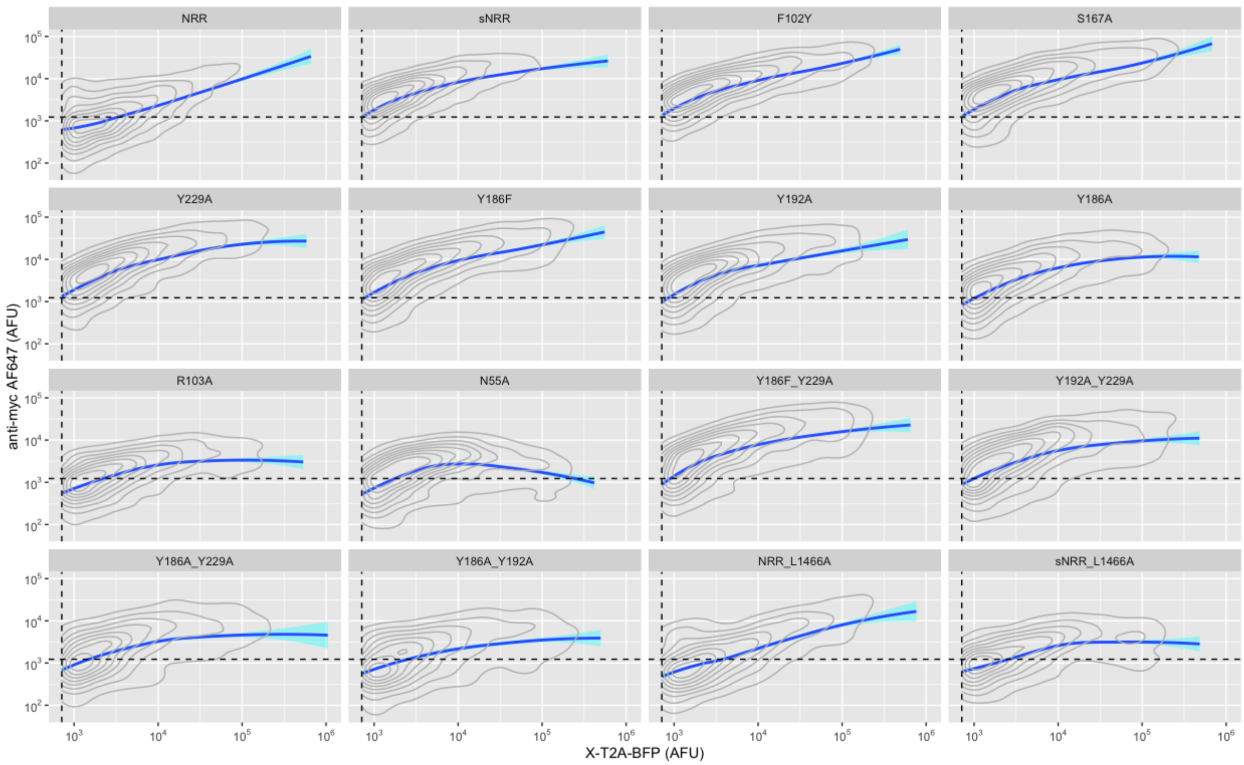


**Fig. S6. Comparison of mechanoreceptors surface presentation levels.** Sixteen individual sNRR domains exhibiting differing mechanical properties were expressed as T2A-BFP fusion constructs via transient transfection in HEK293-FT cells. The surface expression levels of each construct was determined via live cell staining with anti-myc-AlexaFluor488 antibody. Surface staining intensities were then compared against BFP emissions using flow cytometry.

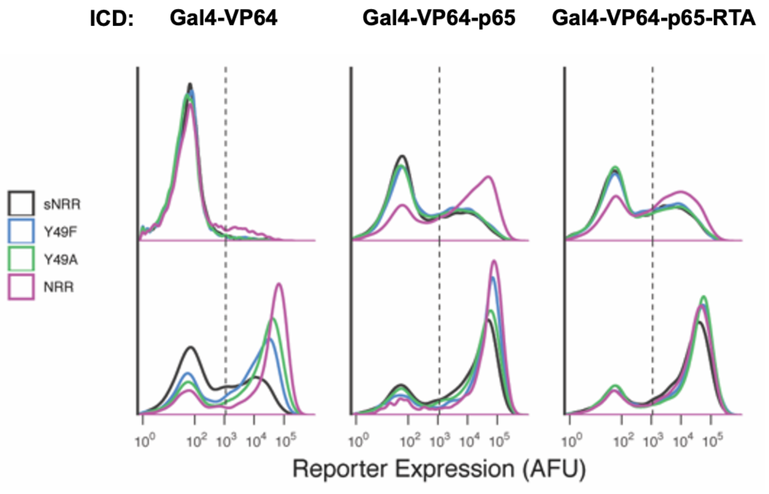


**Fig. S7. Ligand independent activation in receptors.** Receptors containing the indicated ICDs were stably integrated into HEK293-FT cells. ICDs of high transcriptional potency, including VP64-p65 and VP64-p65-RTA, resulted in elevated levels of ligand-independent transcriptional activity due to the strong gene expression mediated by ligand-independently cleaved ICDs. sNRR-based receptors exhibited reduced ligand-independent activity (LIA) compared to those containing NRR.

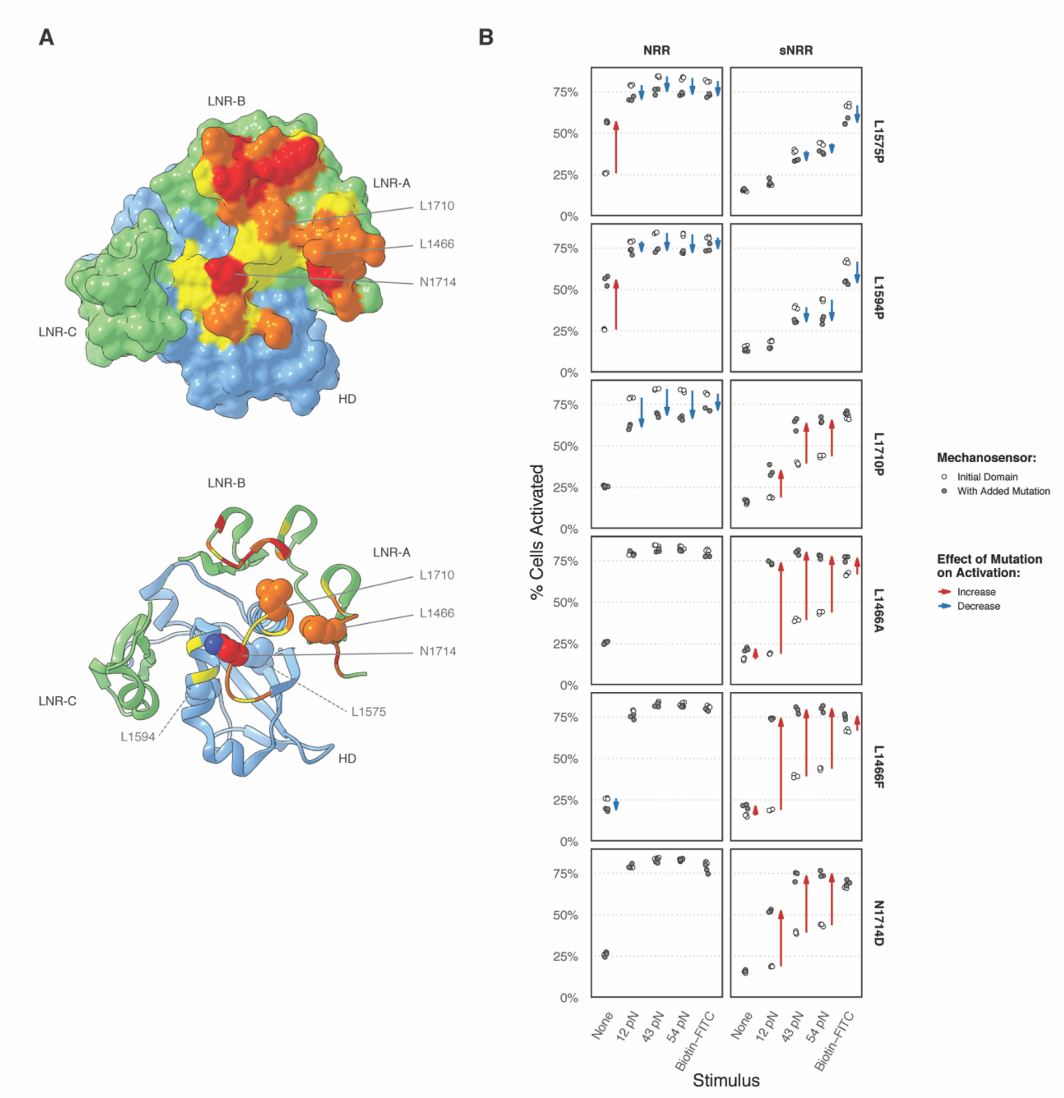


**Fig. S8. Effect of NRR mutations on sNRR tensile strength.** (**A**) Surface view (top) of the Notch1 NRR, with anti-Notch1 Fab binding interface colored in yellow/orange/red, as in main Figure 2B. Mutated residues are denoted. Ribbon view (bottom) reveals all mutated residues in sphere representation, including those internal to the NRR distal to the binding interface (L1575 and L1594). L1575P and L1594P are cancerous mutations, known to destabilize the NRR and distal to the scFv-binding site. L1710P is a cancerous NRR mutation within the scFv-binding interface (PDB 3L95). (**B**) The effect of mutations on receptor strength. Each individual plot depicts the effect of an added mutation (gray circles on plots, vertical axis of grid) on an NRR or sNRR mechanosensing domain (open circles on plots, horizontal axis of grid). Arrows denote the change in mean activation upon mutation, for changes of magnitude > 5%. Mutations that weaken the receptor are expected to increase activation (red), while mutations that strengthen the receptor are expected to reduce activation (blue). L1575P and L1594P destabilize the NRR's autoinhibitory conformation, causing leaky activation in the absence of tension, while not greatly affecting sNRR domains. L1466A, L1466F, and N1714D, which reside at the scFv:NRR binding interface but away from the LNR stabilizing interface in WT NRR, destabilize sNRR domains without affecting NRR activation. L1710P exists at the overlap of NRR's natural stabilizing interface and sNRR's engineered scFv interface, and in turn it impacts signaling of both receptors.

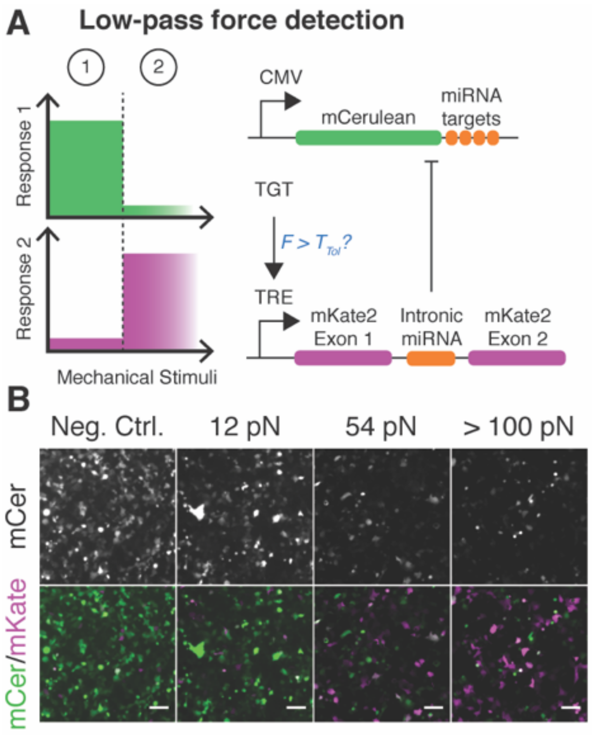


**Fig S9. Low-pass filtering in mechanogenetic circuits.** (**A**) Anticipated signaling responses and circuit diagram showing a constitutively transcribed mCerulean gene, the mRNA transcript of which is targeted for miRNA-mediated degradation via fusion with 3' miRNA targeting sequences. Expression of a corresponding FF4 miR from an intronic TRE-redulated mKate2 construct results in degradation of the mCerulean target mRNAs. Receptors with a tTA ICD induce TRE-driven mKate2 mRNA simultaneously with FF4-miRNA via intronic splicing. TRE-promoter activation thus results in increased mKate2 emission, alongside reduced mCerulean expression, due to miRNA targeting of the mCerulean gene. (**B**) Fluorescence images of HEK293FT cells expressing the low-pass circuit components and stimulated with TGTs. mCerulean expression is only detected only in response to low Ttol TGTs; high $T_{tol}$ TGTs induce mKate2 expression concurrently with reduction of mCerulean. Scale bar 100 µm.

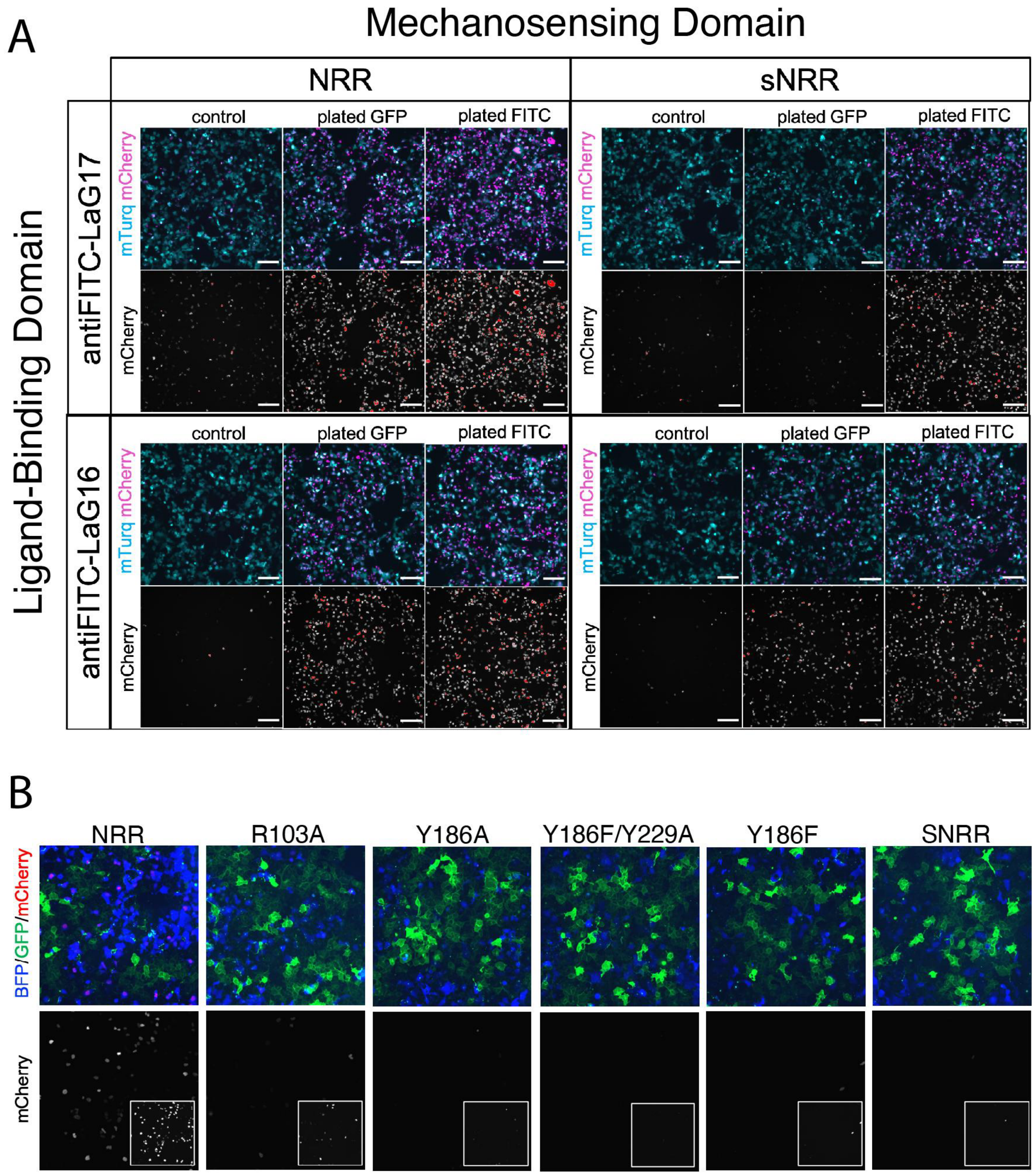


**Fig. S10. LaG16-mediated GFP ligand recognition facilitates sNRR activation.** (**A**) Cells expressing LaG16-sNRR exhibit signaling responses when grown on surfaces containing neutraviding-immobilized mono-biotinylated-GFP, whereas cells-expressing LaG17-sNRR are refractory against the immobilized ligand. Images represent reporter (H2B-mCherry; red) expression in transiently transfected HEK293-FT reporter cells (UAS-H2B-mCherry). A plasmid encoding constitutively expressed by mTurquoise2 was utilized as a transfection marker. Scale bar - 200 um. (**B**) Transduced reporter cells expressing receptors expressing LaG17-based ECDs displayed reduced, or non-detectable signaling activities when grown in coculture with HEK293FT GFP-DLL1 sender cells. These results are in contrast to the trans-cellular action of receivers expressing LaG16-based receptors (displayed in main Fig. 4B). We note that these data suggest a restriction in the propagation of tension across single LaG17-GFP bonds.

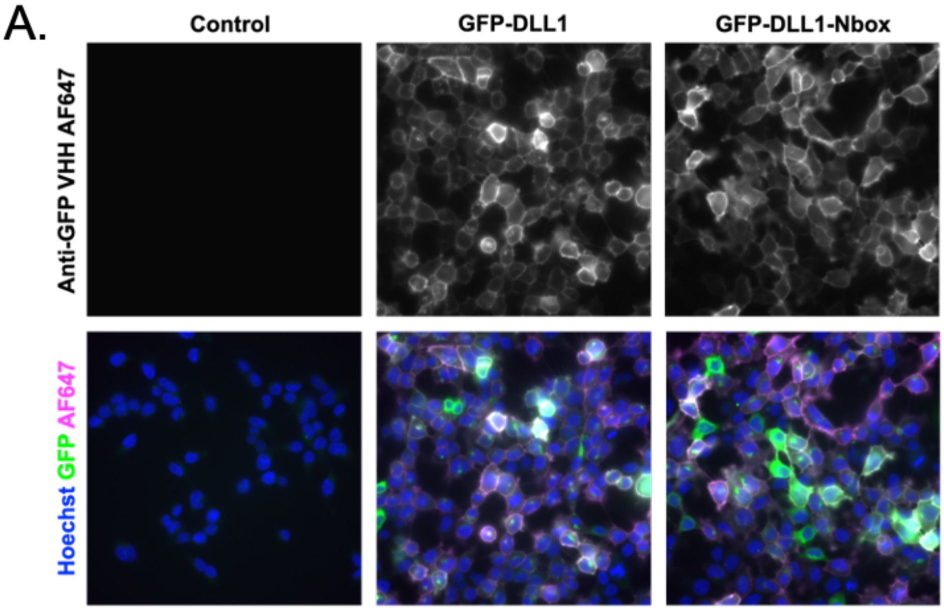


**Fig. S11. Cell surface detection of GFP-based ligands.** (**A**) Transduced sender HEK293-FT cells bearing TRE-regulated GFP-TMD-DLL1 and GFP-TMD-DLL1-Nbox constructs were treated with 100 ng/mL doxycycline for 24 hours prior ro live cell staining with an anti-GFP VHH coupled to AlexaFluor647 ("GFP-Booster AlexaFluor647," AF647; grayscale in top row; magenta in bottom row). Non-transduced HEK293-FT cells were utilized as a control (right-side images); cells were counterstained with Hoescht 33342 to highlight nuclei (blue); direct GFP fluorescence detection is shown in green. Images were collected under live conditions in PBS, following washing out of unbound VHH (see Materials and Methods).

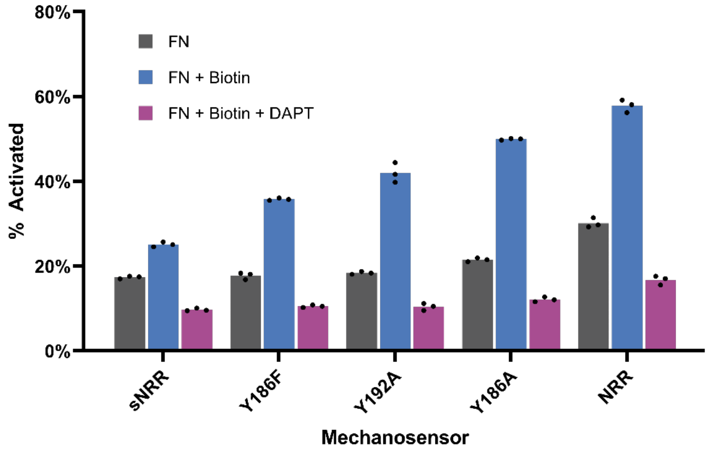


**Fig. S12. Tensile stability of anti-biotin:biotin bonds are sufficient for strengthened receptor activation.** (**A**) The ability of anti-biotin:biotin bonds in mediating sNRR activation was determined by transient transfection of HEK293-FT reporter cells (UAS-H2B-mCherry) with receptors constructs containing anti-biotin-scFv based ECDs. Cells were grown in the presence of wells coated with biotinylated BSA, or in control wells containing coated only with fibronectin. Signaling responses were quantified via flow cytometry, with comparison of cells on control surfaces (gray bars), or on biotin-coated surfaces (blue bars), or in the presence of biotinylated surfaces with gamma secretase inhibition (DAPT at 10 uM). The results confirm the ability of scFv-biotin bonds to withstand the tensile forces required for sNRR activation. Biological triplicate, 2-way ANOVA statistical test show $f < 0.001$ for each between each condition.

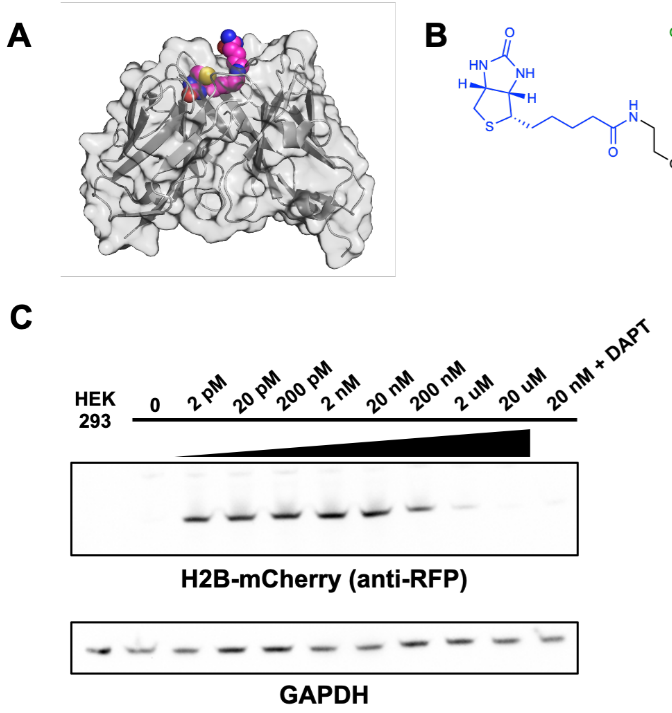


**Fig. S13. Inducible *trans*-cellular coupling using a bifunctional small molecule based on biotin-FITC.** (**A**) Structure of the biotinamide-binding antibody fragment with biotinamide shown in spheres (PDB: 4S1D). (**B**) Chemical structure of the biotin-linked fluorescence (biotin-FITC) compound used. Biotinamide and FITC moieties are shown in blue and green, respectively. (**C**) Western blow of lysates from sender-receiver cocultures. Reporter HEK293-FT (UAS-H2B-mCherry) cells stably expressing anti-FITC-NRR-SynNotch were cultured in combination with stable HEK293-FT senders expressing anti-biotin-TMD-DLL1 ligand. Cells were treated with the indicated concentrations of biotin-FITC, with 20 nM biotin-FITC in combination with 10 uM DAPT or left untreated. Control lysate was derived from non-transfected HEK293-FT cells. Biotin-FITC induced H2B-mCherry expression was probed using anti-RFP antibody (see Materials and Methods). Cells were mixed at 1:1 a sender-receiver ratio and distributed in fibronectin coated 96 well plate microwells. Biotin-FITC was added following adherence (~4 hours later) under conditions of ~95% confluence. Cell lysis was carried out after 24 hours of drug treatment.

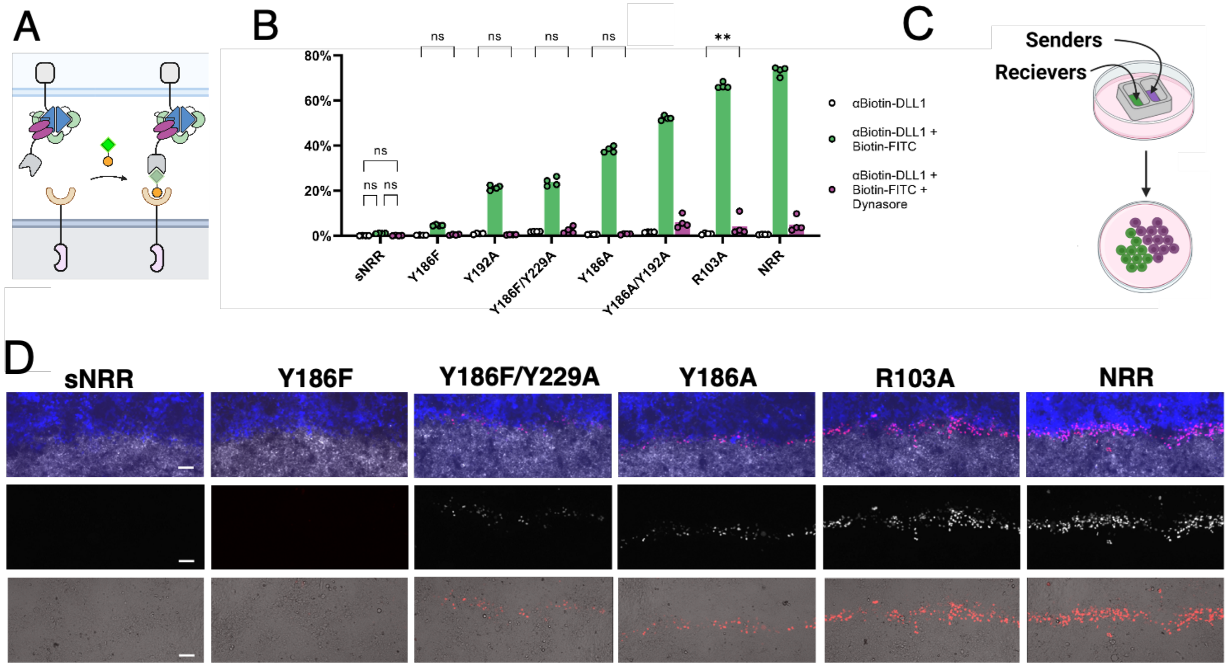


**Fig. S14. Intercellular mechanotransduction via cell-generated endocytic forces.** (**A**) Schematic depicting drug-inducible receptor *trans*-activation. Anti-biotin-TMD-DLL1 ligand cells (bottom) force trans-cellular complexes with anti-FITC receiver cells (top) upon exposure to the upon treatment with the bispecific small molecule bridge biotin-FITC. (**B**) U2OS receiver cell signaling activities quantified from non-patterned sender-receiver mixtures following overnight exposure to biotin-FITC (2 nM; green bars), or following dual treatment with biotin-FITC (2 nM) and dynasore (80 uM; magenta bars). Green and magenta bars represent U2OS receiver cell mixtures with HEK293-FT anti-biotin-TMD-DLL1 senders. Gray bars represent control cocultures in which receivers were mixed with non-ligand expressing HEK293-FT and treated with biotin-FITC (2 nM). Quantifications represent detection of fluorescence reporter levels in U2OS cells bearing an integrated reporter based on UAS-DsRed-Express2, shown as percent activation levels of DsRed+ cells in receptor expressing cells population (T2A-BFP+). Four biological replicates; 2-way ANOVA where null hypothesis cannot be rejected labeled n.s. ($f > 0.05$) ** $f < 0.01$, $f < 0.001$ for all unlabelled comparisons. (**C**) Schematic showing microwell insertion strategy for generating spatially defined sender-receiver interfaces. (**D**) Fluorescence imaging H2B-mCherry reporter expression levels at the interface between sender and receiver populations. HEK293-FT anti-biotin-TMD-DLL1 senders were used in combination with HEK293-FT receivers containing a UAS-H2B-mCherry reporter gene. Cells were treated with 2 nM biotin-FITC 24 hours before imaging. Top images: blue represents receiver cells (receptor-T2A-BFP), white marks sender cells (SNAP-Surface-AlexaFluor-647), red represents H2B-mCherry. Middle and bottom images show H2B-mCherry levels as grayscale (middle), or in red as overlaid with transmitted light (bottom). Scale bars 100 µm.

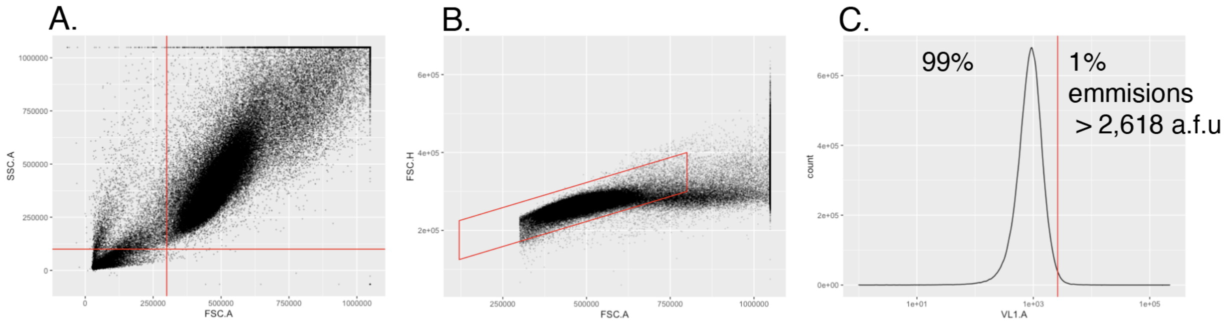


**Fig S15. Flow cytometry gating scheme.** (**A**) Live cells were gated using SSC-A and FSC-A thresholds, defined by the indicated red lines. (**B**) Singlets were isolated using a polygon gate based on FSC-H vs FSC-A as indicated by the red outline. (**C**) Fluorescence gate for isolating receptor expressing cells (in this example T2A-BFP+) was defined as the 99% percentile of control non-transduced cells, analyzed under identical excitation and detection parameters. Same was done for defining reporter activation thresholds.
